# *ztf-16* is a novel heterochronic modulator that opposes adult cell fate in dauer and continuous life histories in *Caenorhabditis elegans*

**DOI:** 10.1101/2022.06.20.496913

**Authors:** Mark A. Hansen, Anuja Dahal, Taylor A. Bernstein, Chani Kohtz, Safiyah Ali, Aric L. Daul, Eric Montoye, Ganesh P. Panzade, Amelia F. Alessi, Stephane Flibotte, Belén Gaete Humada, Marcus L. Vargas, Jacob Bourgeois, Campbell Brown, John K. Kim, Ann E. Rougvie, Anna Zinovyeva, Xantha Karp

## Abstract

Animal development is a complex yet robust process that can withstand lengthy and variable interruptions. In *Caenorhabditis elegans,* adverse conditions can trigger entry into dauer diapause, a stress-resistant, developmentally arrested stage that occurs midway through larval development. Favorable conditions promote recovery from dauer and the completion of development. During larval development, hypodermal seam cells are multipotent and divide at each stage. At adulthood, seam cells differentiate and express the adult-specific COL-19 collagen. The larval-to-adult transition is controlled by a network of genes called the heterochronic pathway, including the LIN-29 transcription factor that directly activates *col-19* expression, and the *let-7* microRNA that indirectly promotes *lin-29* expression. Notably, most heterochronic genes that oppose adult cell fate are dispensable after dauer. We performed a genetic screen for heterochronic genes that act after dauer and identified *ztf-16,* encoding a zinc finger transcription factor in the *hunchback/Ikaros-*like family. We found that *ztf-16* opposes precocious expression of the adult cell fate marker *col-19p::gfp* equally during both life histories, making *ztf-16(-)* the first precocious heterochronic mutant to be unaffected by dauer. Our data indicate that *ztf-16* regulates *col-19p::gfp* via a novel, *lin-29-*independent mechanism. Endogenous *ztf-16b::gfp* expression is regulated by *let-7* and *ztf-16* acts genetically downstream of *let-7,* but *lin-29* is not required for *col-19p::gfp* expression in *ztf-16* mutant larvae or adults. Taken together, this work illuminates a novel aspect of the heterochronic pathway relevant to both dauer and non-dauer development.

## Introduction

In nature, the complex process of animal development occurs in the context of varying environmental conditions. Adverse conditions can lead to a temporary developmental arrest or diapause in animal species from nematodes to vertebrates (Hand et al. 2016). During diapause, juvenile animals are stress-resistant and can withstand conditions otherwise detrimental to their survival. Once conditions improve, animals can exit diapause and resume development. Thus, animals can develop through one of two life histories: one in which development is continuous or one in which development is interrupted by diapause (Hand et al. 2016; Karp 2021). Developmental outcomes are similar in diapause and non-diapause life histories; however, the impact that diapause has on developmental pathways is poorly understood (Karp 2021).

In *Caenorhabditis elegans,* if environmental conditions are favorable early in development, the nematode develops from embryo to adult by progressing continuously through four larval stages (L1-L4) to adulthood (Byerly et al. 1976). Alternatively, adverse environmental conditions sensed early in larval development can trigger entry into a stress-resistant diapause termed dauer (Cassada and Russell 1975). Dauer entry occurs midway through development, after the second larval molt. If favorable conditions are again encountered, the dauer larvae recover and resume development through post-dauer (PD) larval stages (Cassada and Russell 1975). Post-dauer adults are morphologically and structurally similar to adults that did not experience dauer diapause; however, post-dauer adults display altered chromatin and gene expression (Hall et al. 2010).

One cell type whose development has been studied during both continuous and post-dauer life history is a bilateral set of hypodermal cells called seam cells (Sulston and Horvitz 1977; Liu and Ambros 1991). A major function of seam cells is to secrete proteins that make up portions of the collagenous cuticle overlying the hypodermis (Page and Johnstone 2007). Seam cells are multipotent progenitor cells during the larval stages where they undergo self-renewing divisions to produce another seam cell as well as a daughter that fuses with the surrounding hyp7 syncytium. The specific pattern and sequence of seam cell divisions varies with larval stage. Upon reaching adulthood, seam cells exit the cell cycle and terminally differentiate (Ambros and Horvitz 1984). Terminal differentiation and adoption of adult seam cell fate involves several events including seam cell fusion, the secretion of an adult-specific cuticular structure called adult alae, and expression of adult-specific collagens such as COL-19 (Sulston and Horvitz 1977; Ambros and Horvitz 1984; Podbilewicz and White 1994; Liu et al. 1995).

The timing of adoption of adult seam cell fate is controlled by the heterochronic gene network (Ambros and Horvitz 1984). In precocious heterochronic mutants, one or more larval cell division programs are skipped, resulting in the expression of adult cell fate while the worm is still reproductively immature. Alternatively, in reiterative mutants, larval programs are repeated, resulting in reproductively mature worms whose seam cells are still expressing larval programs (Ambros and Horvitz 1984). The most downstream gene in the heterochronic network is *lin-29* (Ambros and Horvitz 1984; Ambros 1989). *lin-29* encodes a transcription factor that most directly promotes the onset of adult cell fate and the characteristics that define it (Rougvie and Ambros 1995; Azzi et al. 2020). In young larvae, *lin-29* expression in the hypodermis is opposed by two other heterochronic genes, *hbl-1,* encoding a hunchback-like transcription factor, and *lin-41,* encoding an RNA-binding protein orthologous to Trim71 in mammals. In particular, LIN-41 represses translation of the *lin-29a* isoform (Slack et al. 2000; Aeschimann et al. 2017; Azzi et al. 2020). In late larval development, *lin-41* expression is silenced by the *let-7* microRNA, thereby allowing expression of *lin-29* and adoption of adult cell fate (Reinhart et al. 2000; Slack et al. 2000; Aeschimann et al. 2019).

Heterochronic genes and seam cell development have been studied primarily during continuous development. In contrast, if larvae enter dauer after the second larval molt, seam cells remain quiescent throughout dauer arrest (Hong et al. 1998). If dauer larvae encounter favorable environments, they will recover from dauer and resume development. Larval seam cell division programs occur in the same pattern and sequence in the dauer and continuous life histories (Liu and Ambros 1991). Interestingly, the genetic mechanisms that underlie this development differ depending on life history. The penetrant phenotypes of many heterochronic mutants are suppressed after dauer, indicating that genes necessary during continuous development are dispensable during post-dauer development. This phenomenon has been observed in both precocious and reiterative mutants (Liu and Ambros 1991; Abrahante et al. 1998; Abrahante et al. 2003; Karp and Ambros 2012). Modulation of microRNA activity after dauer is one mechanism that accounts for the suppression of some reiterative mutants (Karp and Ambros 2012). However, this mechanism does not explain how the loss of function of genes promoting larval programs (precocious mutants) can be tolerated in post-dauer animals.

In order to understand which genes are important for preventing precocious adult cell fate after dauer, we performed a forward genetic screen for mutants that express the *col-19p::gfp* adult cell fate marker as post-dauer larvae. We identified *ztf-16,* which encodes a C_2_H_2_ transcription factor within the *hunchback/Ikaros-*like family (Large and Mathies 2010). *ztf-16* mutants expressed precocious *col-19p::gfp* at all pre- and post-dauer larval stages, as well as during continuous development, making *ztf-16* the first heterochronic gene equally important for opposing adult cell fate during both life histories. We found that ZTF-16 was expressed in a variety of cell types, including the lateral and ventral hypodermis and that this hypodermal expression was cyclical, peaking during the molts. We used genetic and molecular experiments to place *ztf-16* in the context of the known heterochronic pathway, demonstrating that *ztf-16* is regulated by *let-7* but does not appear to be a direct target of *let-7*-induced silencing. Surprisingly, *ztf-16* appeared to act in parallel to *lin-29,* suggesting that *ztf-16* regulates adult cell fate via a novel mechanism. Our results suggest that *ztf-16* acts as a heterochronic modulator and functions primarily to regulate the expression of *col-19*.

## Materials and Methods

### Nematode maintenance

Strains were grown on nematode growth medium (NGM) plates using *Escherichia coli* OP50 as a food source (Brenner 1974). All strains were maintained at 20°C unless they contained the temperature-sensitive *let-7(n2853)* mutation, in which case they were maintained at 15°C. The genotype of all strains in this study is listed in Table 1. Genotypes of newly created strains were confirmed by PCR genotyping and/or sequencing. Table S1 contains a list of genotyping primers.

**Table 1.**
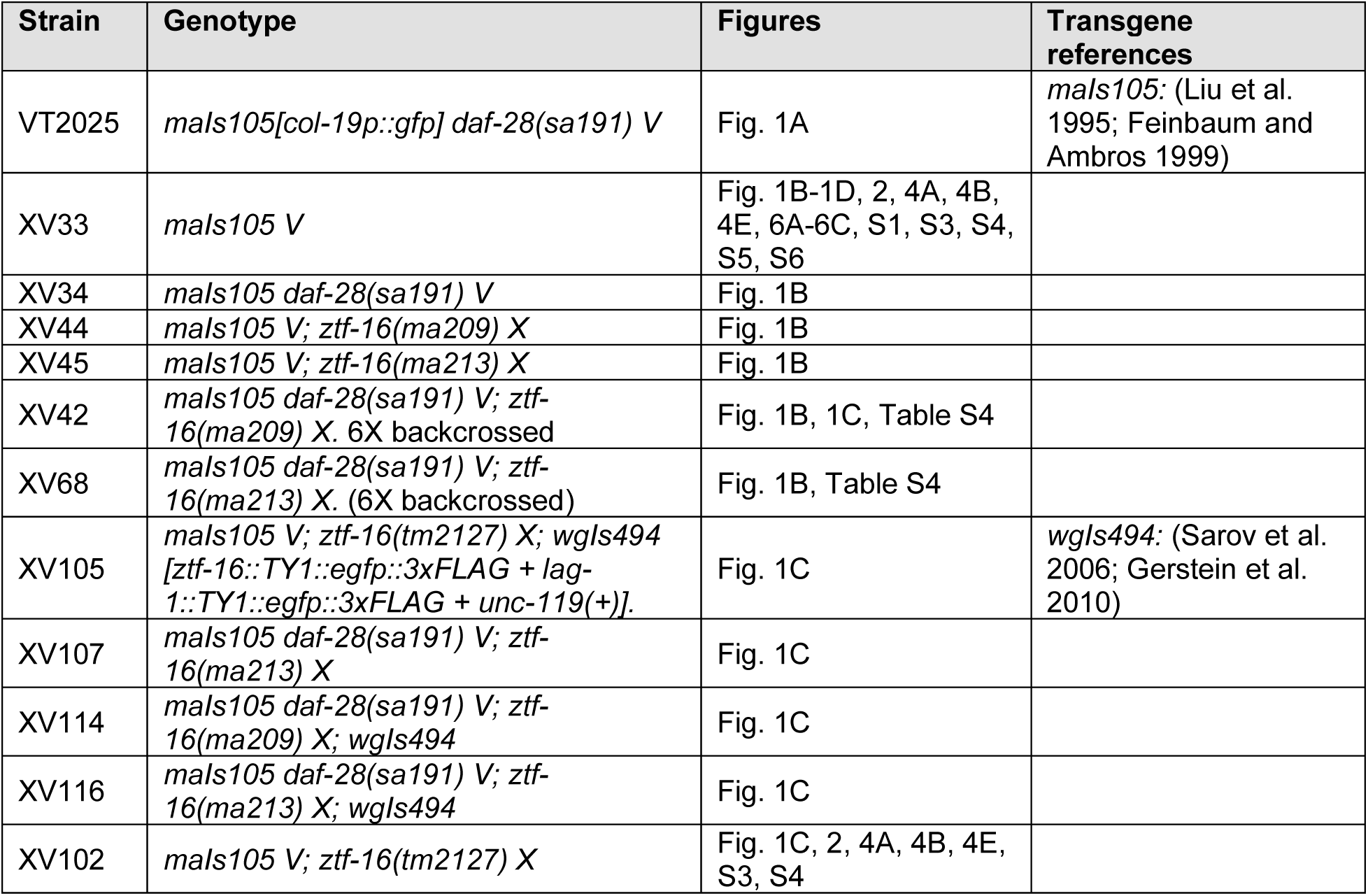
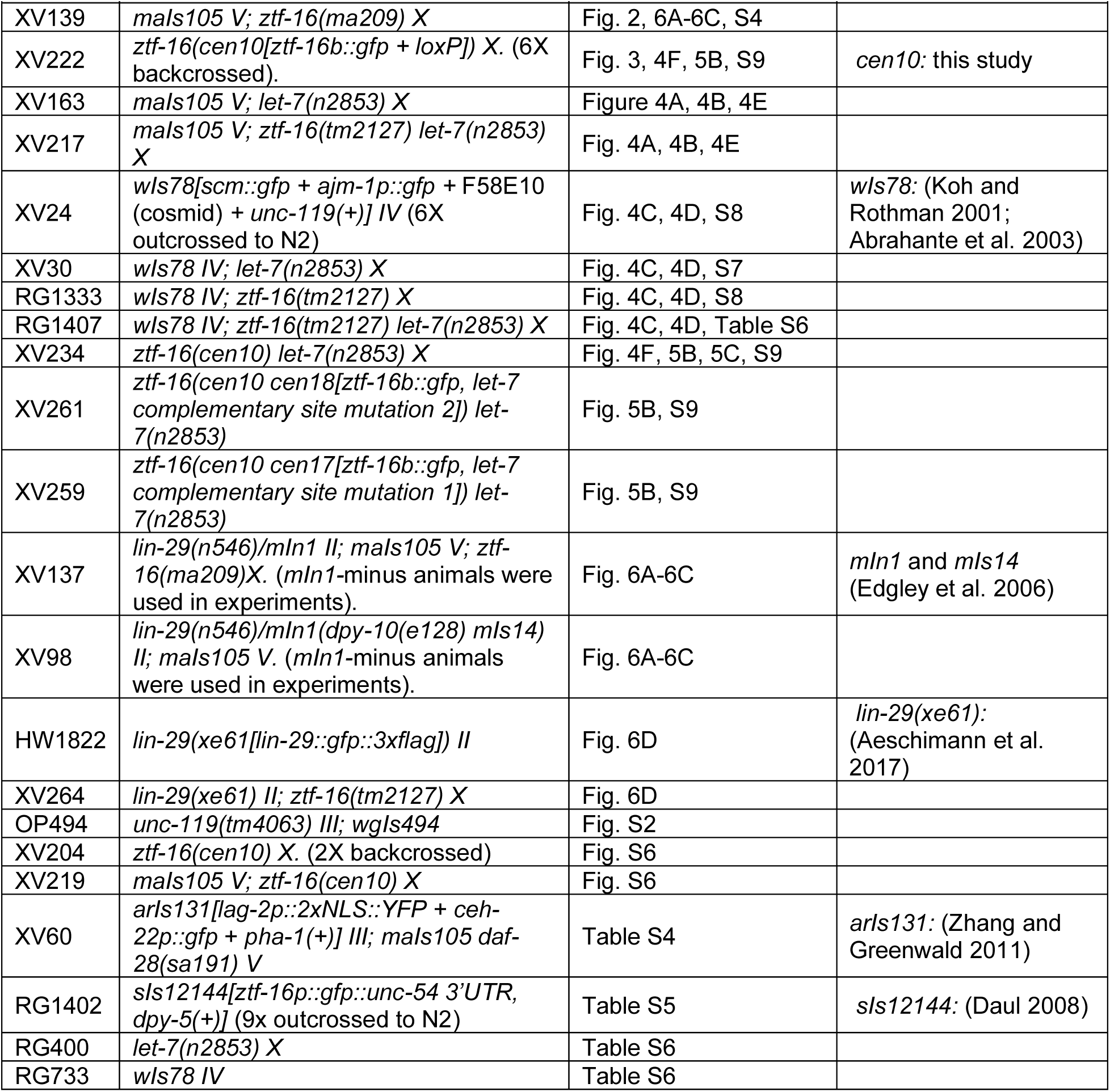
Strains used in this study.

### Genetic screen

L4 staged larvae of the genotype *maIs105[col-19p::gfp] daf-28(sa191)* (VT2025) were incubated in M9 buffer with 50mM ethyl methanesulfonate (EMS) for four hours at 20°C. The larvae were washed and then singled out onto 150 mm plates. After 3 days, when the oldest F1 progeny had developed to the L3/L4 larval stage, they were shifted to 25°C. Two days later the F2 generation had entered dauer due to the *daf-28* mutation. Dauer larvae were selected by treatment with 1% (w/v) SDS and then shifted back to 20°C to induce recovery. Twenty hours later, post-dauer L4 larvae were screened for precocious *col-19p::gfp* expression using a fluorescence dissecting microscope. GFP-positive larvae were singled onto fresh plates. *ma209* and *ma213* were isolated from separate P0 plates. These mutants were both fertile and produced precocious progeny. The isolated alleles were backcrossed 6-8 times, selecting for precocious *col-19p::gfp* expression every two generations.

### Dominance testing

Dominance testing was performed by crossing males heterozygous for a dominant marker *arIs131[lag-2p::yfp, ceh-22p::gfp]* to hermaphrodites homozygous for the relevant *ztf-16* allele. Both males and hermaphrodites were also homozygous for *daf-28* and *maIs105.* Specifically, XV34 *daf-28 maIs105* males were crossed to XV60 *arIs131; daf-28 maIs105.* Male progeny from this cross were then mated to hermaphrodites of the genotypes *daf-28 maIs105; ma209* or *daf-28 maIs105; ma213.* Hermaphrodite cross progeny at the L4 stage that displayed pharyngeal GFP were examined for precocious *col-19p::gfp* using a 1X objective on a fluorescence dissecting microscope (Zeiss Stereo V12 fitted with M2 Bio for fluorescence).

### Whole genome sequencing

Genomic DNA was extracted using the Qiagen Gentra Puregene Tissue Kit (cat. 158667 and 158689) with the following protocol modifications: samples in cell lysis solution were snap frozen, thawed, and vortexed for 30 sec three times, samples were nutated during Proteinase K and RNase A digestions, isopropanol precipitation was supplemented with 3 μL of glycogen and performed overnight at -80°C. The DNA library was created by fragmenting 200 ng of DNA using a Covaris E210. End repair and adapter ligation were then performed using Agilent reagents and an optimized library preparation protocol. Ligated fragments were size selected by purification using SPRI beads while PCR amplification was used to prepare the libraries. The Caliper LabChip GX was used for quality control of the libraries to ensure adequate concentration and appropriate fragment size. DNA sequencing was performed on an Illumina HiSeq 2500 using standard protocols for a 100 bp paired-end run and reads were processed through Illumina’s Real-Time Analysis software. Base calls and their quality scores were generated and CIDRSeqSuite 7.1.0 was used to convert compressed bcl files into compressed fastq files. Paired sequence reads were mapped to the *C. elegans* reference genome version WS230 (www.wormbase.org) using the short-read aligner BWA (Li and Durbin 2010). Single-nucleotide variants (SNVs) were identified and filtered with the help of the SAMtools toolbox (Li et al. 2009). Variant calls also present in the unmutagenized starting strain were eliminated, each SNV was annotated with a custom Perl script, and gene information downloaded from WormBase version WS230.

### Creation of an endogenously tagged fluorescent reporter

Gene editing was done using a combination of the Dickinson method and the SapTrap method (Dickinson *et al*. 2015; Schwartz and Jorgensen 2016). Worms were injected with two guide plasmids, pZ16BsR1 and pZ16BsR2 (see Table S2 and legend), at 25ng/µL each, the pDD162 Cas9 plasmid at 50ng/µL, the repair template plasmid pMLV416 at 50ng/µL, and the *ceh-22p::gfp* injection marker plasmid pCW2.1 at 10ng/µL. The F1 generation was treated with Hygromycin (3 mg/mL) two days later, then screened for rolling worms seven days after injections. Rolling worms were isolated and screened for all-rolling progeny. Positive hits were PCR genotyped to confirm the insertion of the repair template (see Table S1 for genotyping primers). The self-excising cassette (SEC) was removed via heat shock at 34°C for 4 hours and strains were sequenced to confirm the correct integration of the fluorescent protein tag. Primers used in this process can be found in Table S2. The resulting strain was backcrossed 6x prior to use in experiments.

### Mutation of let-7 complementary site

The *let-7* complementary site within *ztf-16::gfp* 3’UTR was mutated using CRISPR/Cas9 genome editing. Commercial *S. pyogenes* Cas9 (IDT, Alt-R® S.p. Cas9 Nuclease V3) was injected at a final concentration of 1.325 µM. The 20µL injection mix also contained 200 µM KCI, 7.5 µM HEPES [pH 7.4], a crRNA targeting *dpy-10* (0.06µg/µL) (Arribere et al. 2014), an Alt-R crRNA (IDT) targeting *ztf-16* 3’UTR (5’-AAAGAAAGUGACAAAUAAAGGUUUUAGAGCUAUGCU-3’, (0.45µg/µL), tracrRNA (0.5 µg/µL), *dpy-10* ss DNA oligo (IDT) (Arribere et al. 2014) (0.1µg/µL), and a single-stranded DNA (ssDNA) oligonucleotides donor (0.2µg/µL) that, when incorporated, replaces the *ztf-16* 3’UTR *let-7* complementary site with an EcoRI restriction digest site. The ssDNA Ultramer DNA oligo (5’-CGGTTTTCTTCGGTTTTTTTCGATGCATATTCACAGTCAATTCTGATGAATTCTTTATTTGTCACTTTCTTTCTTCATAACAGGGAAACAACGAATG-3’) was purchased from IDT. The following primers were used to amplify the region surrounding the *let-7* complementary site (F: 5’-CCCCATTCTTTCATCCACAT-3’; R: 5’-GGTGGTGGTGAGAAAGGAAA-3’); an EcoRI restriction digest of the resulting PCR fragment positively identified a successful edit. Two *let-7* complementary site mutations *ztf-16(cen17)* and *ztf-16(cen18)* were confirmed by sequencing three independent lines.

### RNAi

The *ztf-16* RNAi clone was pHG49 (Gardner 2005). The *lacZ* RNAi clone was pXK10 (Karp and Greenwald 2003). The *lin-41* clone was from the Ahringer library (Source Bioscience) (Kamath et al. 2003). RNAi plates were prepared by adding 200µL of an overnight culture of RNAi bacteria to 60mm NGM plates containing 50µg/mL carbenicillin and 1mM IPTG. Embryos were then isolated from a population of gravid adults by two successive two-minute treatments with a solution of 10% (v/v) Clorox bleach in 1 M NaOH and added to the RNAi plates. Worms were incubated at 24°C until the L4 stage and then scored as described below.

### Obtaining synchronous populations

Synchronous populations of continuous development stages were obtained by allowing gravid adult hermaphrodites to lay eggs at 24°C on a seeded 60 mm plate for 2-4 hours. Experiments involving *let-7* mutants which do not lay eggs well were isolated by bleaching as described above. The embryos were incubated at 24°C until they reached the desired stages. Table S3 lists the incubation times used for each stage. Precise stages were confirmed by gonad morphology using DIC microscopy. Only worms of the desired stage were scored. Prior to mounting, pharyngeal pumping was monitored to identify molting (lack of pumping) or intermolt (pumping) stages. Strains examined during continuous development were wild-type for *daf-28.* Pre-dauer L1 and L2d-staged larvae were obtained in a similar manner to L1 and L2 staged larvae during continuous development, except that all strains contained *daf-28(sa191)*.

Dauer formation was induced by allowing worms to crowd and starve at 15°C for *let-7(n2853)* experiments, or at 24°C for all other experiments. Dauer larvae were selected by treatment with 1% (w/v) SDS at room temperature for 30 minutes, except for the *ztf-16b::gfp* experiments in Fig. 3, where dauer larvae were identified by morphology, including dauer alae (Cassada and Russell 1975; Karp 2018). Post-dauer populations were obtained by SDS-selecting dauer larvae from starved plates as described above and then returning dauer larvae to 24°C to recover. Table S3 lists the incubation times used to reach different stages. Experiments with strains containing wIs78 were performed at 25°C.

### Microscopy and phenotypic characterization

Worms were picked to a 2% agarose pad in 10 mM levamisole. All phenotypes were scored using a Zeiss Axio Imager D2 with an AxioCam MRm camera and a Zeiss Illuminator HXP 200 light source. GFP expressed by *maIs105 [col-19p::gfp]* was scored using a 40X objective with an exposure between 8-30 msec (see Figure legends for each experiment). For each worm, three images were taken of the hypodermis, one of the anterior, one centred on the vulva, and one of the posterior. Based on these images, detectable expression was placed into one of three categories: bright, moderate, and dim. Representative images for these categories are shown in Fig. S1. *wIs78* [*ajm-1::gfp* + *scm::gfp*] expression was scored using a 63X objective with an exposure of 150ms. *ztf-16(cen10*[*ztf-16::gfp* + *LoxP*]) expression was scored using a 40X objective with an exposure time of 250ms. *lin-29::gfp* expression levels were examined with *xe61[lin-29::gfp::3x-flag]* (Aeschimann et al. 2017) using a 40X objective with an exposure time of 372.7 ms. *wgIs494* [*ztf-16::TY1::egfp::3xFLAG; unc-119(+)*] expression was scored using a 40X objective with an exposure time of 250ms. For comparing *ztf-16(cen10[ztf-16::gfp])* expression levels, a 63X objective was used with an exposure time of 10-250ms (see Figure legends for details). To better visualize the dim images presented in Figure 3B, the Lighten Shadows function in Adobe Photoshop Elements 14 was adjusted by +1%. These adjustments were made across the entire image and for all images shown in this Figure.

For comparing expression when fluorescence levels appeared similar, we used ImageJ. GFP-positive nuclei were outlined using the freehand tool and the mean pixel intensity for all of such nuclei were averaged together for each individual worm. Adult alae were scored using DIC optics and a 63X objective. The worms scored for alae were the same worms scored for *col-19p::gfp*.

### mRNA-sequencing

Strains XV102 *col-19p::gfp; ztf-16(tm2127),* XV139 *col-19p::gfp; ztf-16(ma209),* and XV33 *col-19p::gfp* were grown in parallel. Post-dauer samples were obtained by allowing strains to crowd and starve at 24°C, then selecting dauer larvae in 1% (w/v) SDS for 30 minutes. Dauer larvae were added to fresh seeded plates and incubated at 24°C for 25-26 hours, at which time the population was in the post-dauer L4 stage. Continuous L4 samples were obtained by sodium hypochlorite treatment of a population of gravid adult hermaphrodites to isolate embryos. Embryos were then grown at 24°C for 40 hours, at which time the population was in the L4 stage. PDL4 or L4 staged larvae were washed with M9 buffer and centrifuged. Pellets of approximately 100µL of packed worms were treated with 1mL of TRIzol reagent (Invitrogen) and frozen in dry ice/ethanol. RNA preparation was performed as previously described (Zinovyeva et al. 2014). Samples were sent to Genewiz for library preparation, poly-A selection, and Illumina HiSeq sequencing (Illumina HiSeq 4000).

### Quality check and filtering of RNA-seq samples

Paired-end (PE) RNAseq reads were obtained from continuous development (CD) and post-dauer (PD) life histories for the following strains: XV102 *col-19p::gfp; ztf-16(tm2127),* XV139 *col-19p::gfp; ztf-16(ma209),* and XV33 *col-19p::gfp*. Initially, three biological replicates were obtained for all strains with the exception of XV102 (CD), which had two biological replicates. The libraries were un-stranded 150bp in length for both reads in pairs. The raw reads were assessed for quality using fastqc (https://www.bioinformatics.babraham.ac.uk/projects/fastqc/) before and after filtering. A thorough analysis was performed for over-represented sequences (such as adapter sequences or PCR “hotspots”). The quality-based filtering was done using trimmomatic tool with the following parameters ‘ILLUMINACLIP: TruSeq3-PE-2. fa: 2:25:25 LEADING:3 TRAILING:5 SLIDINGWINDOW:4:15 MINLEN:18’ (Bogler *et el.* 2014). The TruSeqv3 kits adapter sequences were used for read trimming for both reads in a pair. The quality score was checked for every base with at least a 4-nucleotide sliding window size per base. The surviving reads in fastq format were given as input for further analysis.

### Genomic alignment and expression quantification

The quality-filtered reads were mapped to the *C. elegans* reference genome (downloaded from Wormbase for version WB279) by STAR aligner with default parameters in two-pass mode to obtain genomic alignment while using reference genome annotation (Dobin et al. 2013). The alignment output was taken into a sorted BAM file for transcripts assembly in a further step. Each genomic alignment BAM file was provided into stringtie transcripts assembly program with “-j 5 -c 5 -B –e” parameters to construct scaffolds and contigs by utilizing genomic annotation from GTF file (Kovaka et al. 2019). This output was given into merge program of stringtie to construct consensus and unique reference transcriptome from all samples. The normalized expression for each gene in the transcriptome was collected from the final output of above step in the form of TPM and FPKM. While we were interested to see differences between the wild-type and mutant samples from CD and PD groups, gene-wise reads count was obtained by utilizing feature Counts program of subread package by providing each sample BAM and reference annotation file while setting a parameter for “gene” as keywords to fetch reads (Yang Liao et al. 2019). The per gene read count from each sample was collected and formed a matrix for differential expression using the DESeq2 package in R bioconductor (Love et al. 2014). We performed some basic cleaning over the read count data by removing low counts genes with less than 5 reads across the samples. We tested for reproducibility among biological replicates through principal component analysis (PCA) grouping. We removed outlier replicates, leaving us with two biological replicates for all samples except XV102 PD and XV139 PD, which had three replicates each. The sample information was provided into design matrix for wild-type and mutant strain samples with replicates. The regularized transformation was done by transforming the counts data into log2 scale to fit the model for each sample using prior distribution of coefficients which was estimated from the data. The differential expression results were obtained by making a contrast between the mutant and wild-type groups. Afterward, the differential expressed genes were counted based on their log2Foldchange and adjusted p-value significance (<= 0.05).

### Geneset enrichement analysis (GSEA) and visualization of important genes

The list of genes according to their differential expression status was used as input for geneset enrichment analysis. We used WebgeStaltR, an R package for overrepresented genes analysis from the provided list (Yuxing Liao et al. 2019). We followed two different criteria to determine significant categories based on their input gene number, 1) a significant FDR value cutoff of <= 0.05 and 2) top categories without considering FDR.

The categorization of some important genes according to their significance was pulled out from the differentially expressed genes such as high-confidence genes, heterochronic genes, etc. using python and R in-house written scripts. All plots were generated using in-house scripted python and R codes.

### Statistical Analysis

For *col-19p::gfp* expression and alae comparisons, the Freeman-Halton extension of the Fisher Exact test (VassarStats) was performed. Seam cell number was compared with the two-tailed *t* test (Graphpad PRISM Version 8). The Fisher Exact test (Graphpad PRISM Version 8) was used to compare the number of apical junctions between seam cells. Mean pixel intensity was compared using the Kruskal-Wallace and Dunn’s test or the Mann Whitney test, as appropriate (Graphpad PRISM Version 9). See Figure legends for details.

### Data Availability

Sequencing data files are available on the NCBI Sequence Read Archive (SRA) under the accession number SUB11568807 with bioproject id PRJNA848055. Most of the tertiary analysis was performed by in-house written custom codes in R and python. All other data necessary for confirming the conclusions presented in the article are represented fully in the article, figures, tables, and supplemental material. All strains and plasmids generated for this study and their associated sequence files are available upon request.

- **Table S1** lists all DNA primers used for PCR genotyping.
- **Table S2** lists the sequences of DNA primers used for CRISPR-editing to tag *ztf-16b* with *gfp*.
- **Table S3** shows the time in hours required for *C. elegans* to develop to the specified stage at 24°C.
- **Table S4** shows the results of dominance testing for *ma209* and *ma213*.
- **Table S5** shows the expression of a *ztf-16p::gfp* transcriptional reporter.
- **Table S6** shows the suppression of the *let-7(-)* bursting phenotype by a *ztf-16* mutation
- **Fig. S1** shows representative images of the three categories *col-19p::gfp* expression (dim, moderate, and bright).
- **Fig. S2** shows the hypodermal expression pattern of *wgIs494 [ztf16::egfp]* from L3 to adult.
- **Fig. S3** shows qPCR data for confirmation of mRNA-seq data.
- **Fig. S4** shows high confidence genes regulated by *ztf-16* during continuous development.
- **Fig. S5** shows differential gene expression in post-dauer L4 vs. continuous L4 in wild-type larvae.
- **Fig. S6** shows that the *ztf-16b::gfp* allele created via CRISPR-mediated gene editing is functional.
- **Fig. S7** shows the re-emergence of junctions between seam cells of young adult *let-7(n2853)* mutant worms at raised at restriction temperatures.
- **Fig. S8** shows that *ztf-16* mutants have small gaps between adjacent seam cells.
- **Fig. S9** shows that *let-7* appears to regulate *ztf-16b::gfp* expression indirectly during continuous development.
- **File S1** provides the complete DEG and GO-term enrichment data referred to in the text and/or figures.

## Results

### *ztf-16* blocks expression of an adult cell fate marker after dauer

Heterochronic genes control the timing of adult cell fate, including expression of the adult cell fate marker *col-19p::gfp* (Liu et al. 1995; Rougvie 2001). Prior genetic screens used alterations in the timing of *col-19p::gfp* expression to identify new heterochronic genes or interactors that act during continuous development (Abrahante et al. 1998; Hayes and Ruvkun 2006). To identify genes that are important for proper seam cell fate progression after dauer, we performed a forward genetic screen for mutants displaying precocious expression of *col-19p::gfp* during the post-dauer L4 (PDL4) stage (Fig. 1A). Dauer formation was induced using the *daf-28(sa191)* mutant which forms dauer larvae at elevated temperatures and recovers synchronously at lower temperatures (Li *et al*. 2003) (see Methods). After mutagenesis with EMS, we screened F2 PDL4s for precocious *col-19p::gfp* expression and obtained *ma209* and *ma213* (Fig. 1A). These alleles were backcrossed 6-8X before analysis.

**Figure 1.**
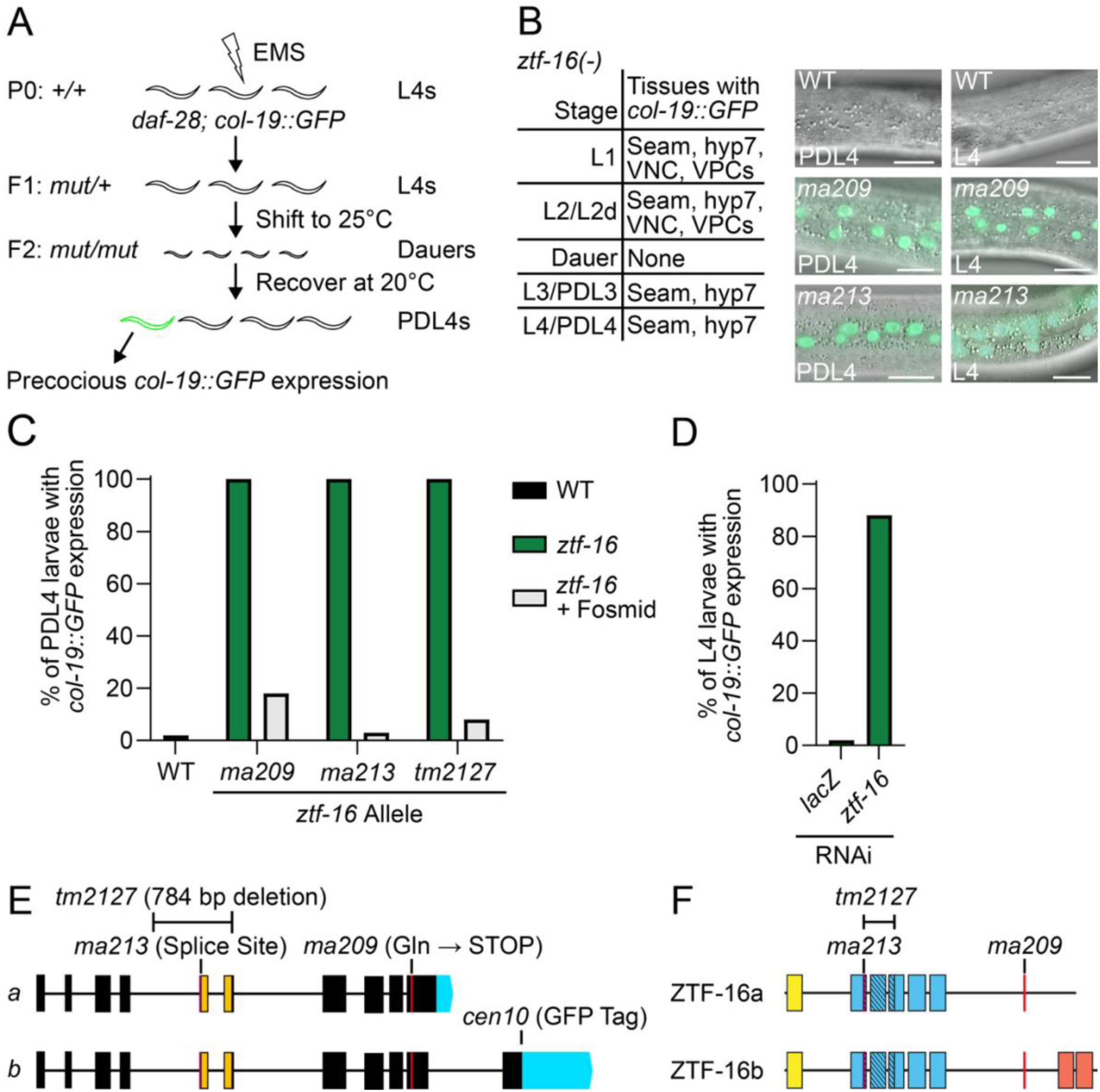
*ztf-16* prevents precocious *col-19p::gfp* expression during post-dauer and continuous development. (A) Schematic of the forward genetic screen to identify post-dauer heterochronic mutants using *col-19p::gfp* expression as an indicator of adult cell fate. (B) *col-19p::gfp* was expressed precociously in pre-dauer, post-dauer, and continuously developing larvae in *ma209* and *ma213* mutants. For worms scored during the dauer stage, dauer formation was induced by starvation/crowding, *daf-28(sa191)*, or both. Post-dauer worms were recovered from *daf-28* populations via SDS selection (n = 17-55 worms/stage/allele). Seam, seam cells; VNC, ventral nerve cord; VPCs, vulval precursor cells. Scale bar = 20μm. (C) Worms displaying any precocious *col-19p::gfp* are shown. *ztf-16* mutants have bright, penetrant *col-19p::gfp* expression in hyp7 and seam cells while *ztf-16* mutants with the fosmid *wgIs494 [ztf-16::egfp]* have dim *col-19p::gfp* limited to the anterior of the worm (n = 12-67 worms/allele). We interpreted any GFP expression as *col-19p::gfp* because *ztf-16::egfp* is not expressed in the lateral hypodermis of wild-type worms during the L4 stage (Figure S2). In addition, the expression of *ztf-16::egfp* during other stages is significantly dimmer than that of *col-19p::gfp*. (D) Worms fed *ztf-16* dsRNA display precocious *col-19p::gfp* expression (n = 50 worms/strain). (E) Schematic of *ztf-16a* and *ztf-16b* isoforms. Boxes represent exons and lines represent introns. Blue shading represents the 3’-UTR. The yellow shading indicates the extent of the exons affected by the *tm2127* deletion. The *cen10* allele is a *gfp* insertion located immediately 5’ of the stop codon of *ztf-16b*. (F) Protein schematic of ZTF-16a and ZTF-16b. Boxes represent zinc-finger domains. Yellow, blue, and red shading represents N-terminal, middle, and C-terminal zinc fingers, respectively. Diagonal hatching indicates zinc fingers removed by the in-frame *tm2127* deletion.

To further characterize the phenotype of *ma209* and *ma213* mutants during the dauer life history, *daf-28; ma209* and *daf-28; ma213* were grown at 24°C to induce dauer formation. Both alleles were recessive (Table S4) and displayed identical phenotypes. Beginning in the L1 stage, we observed *col-19p::gfp* expression in the hypodermis, including seam cells and hyp7 nuclei (Fig. 1B). Hypodermal *col-19p::gfp* expression was evident during the L2d stage, but absent from dauer larvae. Dauer larvae lacked *col-19p::gfp* expression whether dauer formation was induced by *daf-28* or by allowing worms to crowd and starve out. Hypodermal *col-19p::gfp* returned in PDL3 and remained strongly expressed in PDL4 (Fig. 1B). Post-dauer adults expressed *col-19p::gfp* at levels similar to wild type. Despite the penetrant expression of this adult cell fate marker, we found that adult alae were not produced precociously. Surprisingly, we also observed *col-19p::gfp* expression in non-hypodermal cell types (Fig. 1B). The ectopic expression of *col-19p::gfp* was limited to early larval stages (L1-L2) and was not further explored.

To ask whether *ztf-16* also blocks *col-19p::gfp* expression during continuous development, we outcrossed *ma209* and *ma213* to remove *daf-28(sa191).* We grew synchronous populations at 24°C for comparison with the experiments described above. We found that *ma209* and *ma213* mutants that developed continuously displayed the same phenotypes as pre- and post-dauer larvae and adults (Fig. 1B). In contrast, the phenotypes of all the previously identified precocious heterochronic mutants that have been tested are affected by life history (Liu and Ambros 1991; Abrahante et al. 1998; Abrahante et al. 2003).

We hypothesized that *ma209* and *ma213* were alleles of the same gene, as they displayed identical phenotypes. Consistent with our hypothesis, both alleles mapped to LGX. To determine the causal mutations, we performed whole-genome sequencing on *ma209* and *ma213* (6-8X backcrossed), as well as the unmutagenized starting strain. We then looked for X-linked genes in which the coding sequence was affected in both *ma209* and *ma213.* A single candidate gene emerged from this analysis: *ztf-16*.

To confirm *ztf-16* as the causal mutation, we first compared our alleles to an existing *ztf-16* deletion allele, *tm2127*. Similar to *ma209* and *ma213, tm2127* mutants displayed precocious *col-19p::gfp* expression but adult alae formation at the same time as wild type in both life histories (Fig. 1C). Furthermore, reduction of *ztf-16* activity via RNAi also caused precocious *col-19p::gfp* expression (Fig. 1D). Finally, we performed rescue experiments using an available transgene (*wgIs494*) containing a fosmid with the *ztf-16* (*egfp*-tagged) locus plus the surrounding genomic sequence, including the two neighboring genes (see Fig. S2 legend). The fosmid efficiently rescued the precocious *col-19p::gfp* expression in *ztf-16* mutants (Fig. 1C).

Prior work established that *ztf-16* regulates gonad morphogenesis and glial cell remodeling, but no effect on adult cell fate has been reported (Large and Mathies 2010; Procko et al. 2012). *ztf-16* encodes a C_2_H_2_ <u>z</u>inc-finger transcription factor within the *hunchback/Ikaros*-like (HIL) family. Other members of the HIL family in worms are encoded by *hbl-1*, another heterochronic gene, and *ehn-3* (Large and Mathies 2010). Members of the HIL family have a particular arrangement of zinc fingers with an N-terminal or central cluster that acts as a DNA-binding domain and a pair of C-terminal fingers that mediate protein-protein interactions (Reviewed in (Iuchi 2001)). *ztf-16* encodes two isoforms: a *ztf-16a* isoform and a larger *ztf-16b* isoform (Figures 1E-F). The *ztf-16b* isoform retains the HIL-family zinc-finger arrangement, while the *ztf-16a* isoform has an abbreviated final exon that lacks the two C-terminal zinc fingers (Fig. 1F). Furthermore, the *ztf-16b* isoform contains a longer 3’UTR than the *ztf-16a* isoform (Fig. 1E). The alleles we obtained from our screen disrupt the zinc finger arrangement of ZTF-16 by producing a premature stop codon (*ma209*) or altering the splicing of the transcript (*ma213*) (Fig. 1E). *tm2127* also disrupts the zinc finger arrangement as three of the central zinc fingers are compromised by the in-frame deletion (Fig. 1E-F).

### Regulation of gene expression by *ztf-16*

To determine how the ZTF-16 transcription factor impacts gene expression, we sequenced mRNA from PDL4 and continuous L4 whole larvae in *ztf-16(tm2127), ztf-16(ma209),* and control larvae. For each life history, we compared gene expression between each mutant and control (File S1). We then identified a high confidence set of genes whose expression changed significantly (FDR ≤ 0.05) in the same direction in both mutants. Using this approach, we found that in PDL4 *ztf-16* mutants, 211 genes were upregulated and 93 genes were downregulated (Fig. 2A). As expected, *col-19* levels were significantly upregulated in *ztf-16(tm2127)* PDL4 larvae, but surprisingly, *col-19* levels were not affected in *ztf-16(ma209)* PDL4 larvae (File S1). However, this analysis was complicated by the presence of the multicopy *col-19p::gfp* transgene in the strains that were sequenced. High confidence genes that were upregulated in post-dauer *ztf-16* mutants were involved in signaling, gland development, and collagens, while the genes that were downregulated in post-dauer *ztf-16* mutants were involved in the immune response (Fig. 2B). To confirm these results, we hand-picked control and *ztf-16(tm2127)* post-dauer larvae at the late L4 stage (L4.5-L4.7) (Mok et al. 2015) and performed qPCR. We found that the expression of 4/6 downregulated genes and 6/10 upregulated genes reproducibly changed in the same direction in the qPCR experiments (Fig. S3).

**Figure 2:**
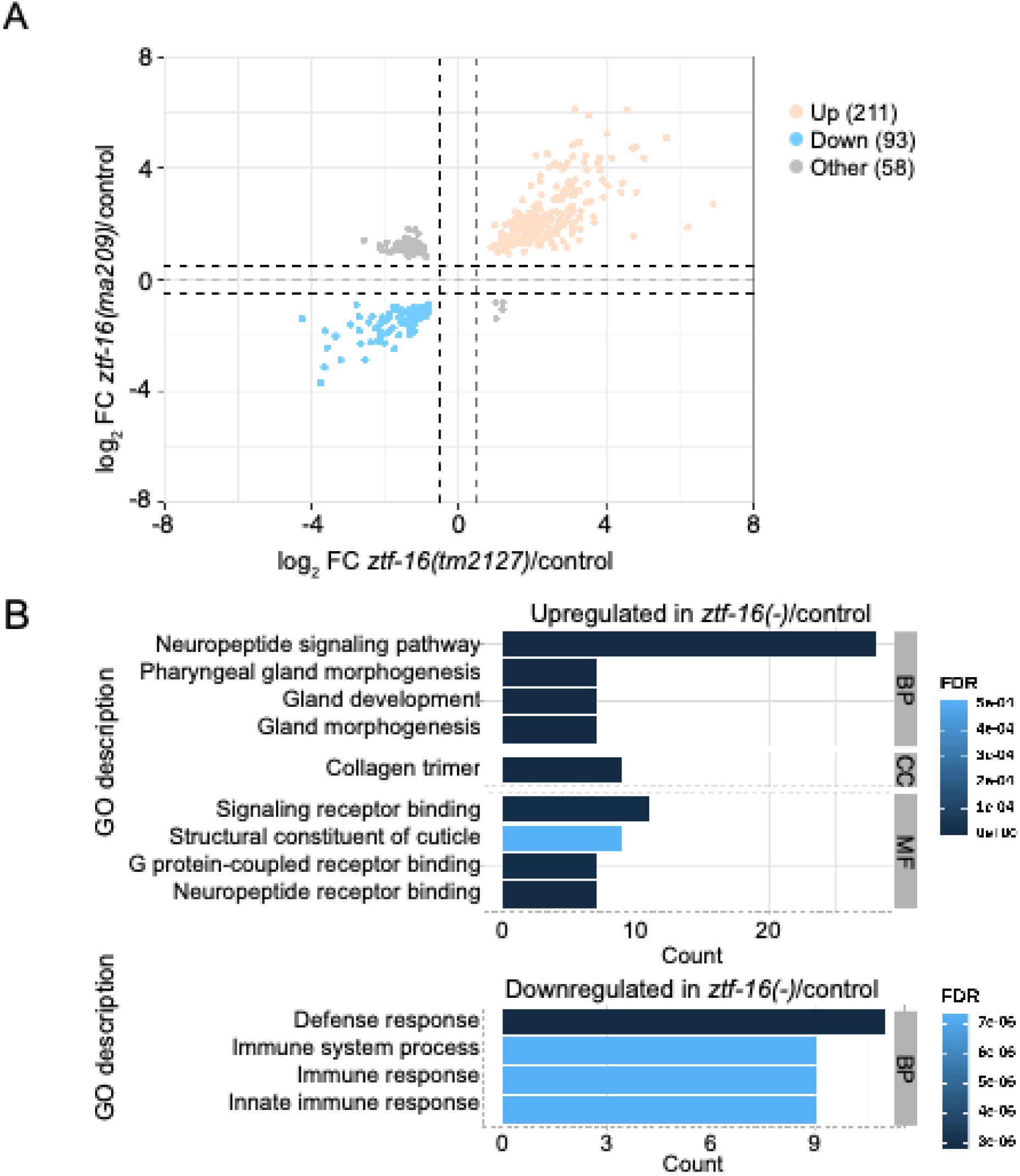
Gene expression changes in *ztf-16* mutant post-dauer larvae. mRNA-seq data were collected from two different *ztf-16* mutants and wild-type controls during the PDL4 stage. (A) A volcano plot showing all genes whose expression changed significantly (FDR ≤ 0.05) in *ztf-16* mutants relative to controls. (B) Significantly enriched GO terms for the upregulated or downregulated genes. A complete list of genes, log2 fold-change, FDR, and enriched GO terms is given in File S1. BP = Biological Process, CC = Cellular Component, MF = Molecular Function, FDR = False Discovery Rate.

In continuous development, 54 genes were upregulated, and 9 genes were downregulated in *ztf-16* mutants (Fig. S4A). The genes that were upregulated in *ztf-16* mutants during continuous development were involved in cuticle development, molting, and collagens (Fig. S4B). There were too few downregulated genes for statistically significant enrichment, however these genes appeared to be involved in responses to environmental cues and cellular stress responses (File S1). Notably, 14/54 (26%) upregulated genes and 3/9 (33%) downregulated genes were also significantly changed in the same direction after dauer, suggesting that these genes are regulated similarly by *ztf-16* in both life histories (File S1).

During the analysis described above, we noticed that approximately three times as many genes were differentially regulated in *ztf-16* mutant PDL4 larvae as in continuous L4 larvae, suggesting that *ztf-16* may be more important to regulate gene expression in the dauer life history (Figs. 2, S4). This observation prompted us to wonder how gene expression was altered in wild-type PDL4 larvae relative to continuous L4. We found that 1003 genes were upregulated and 782 genes were downregulated in post-dauer larvae (log_2_ fold change ≥1 or ≤ -1, FDR ≤ 0.05) (Fig. S5). Using GO-term analysis, we found that downregulated genes were involved primarily in metabolic processes, and upregulated genes were involved in stress response, immune response, cell cycle, and molting (File S1).

### Endogenous *ztf-16b::gfp* expression oscillates with the molting cycle

Heterochronic genes act at particular developmental timepoints, and heterochronic gene expression is typically temporally regulated (Rougvie 2001). In order to view when and where ZTF-16 is expressed, we used CRISPR-Cas9 to introduce a C-terminal GFP tag, producing the *cen10[ztf-16b::gfp]* allele (Fig. 1E) (Dickinson *et al*. 2015; Schwartz and Jorgensen 2016). We focused on the *ztf-16b* isoform because that isoform includes the C-terminal zinc fingers. Tagged ZTF-16 is functional because *ztf-16(cen10)* blocked precocious *col-19p::gfp* expression nearly as effectively as wild-type *ztf-16* (Fig. S6). We were particularly interested in the lateral hypodermis, as this is the primary tissue in which *col-19p::gfp* is precociously expressed. Hypodermal expression of a C-terminal *ztf-16b* reporter transgene was previously mentioned, though the focus of that work was the somatic gonad (Large and Mathies 2010). We found that endogenous *ztf-16b::gfp* was expressed in the lateral hypodermis, including seam cells and hyp7 nuclei (Fig. 3). We also observed *ztf-16b::gfp* in hyp6 and the ventral hypodermis, as well as two non-hypodermal cell types: the distal tip cells and the ventral nerve cord (Fig. 3A). Consistent with prior reports, *ztf-16b::gfp* was localized to the nucleus in all of the tissues in which expression was seen (Procko et al. 2012). Temporally, *ztf-16b::gfp* expression oscillated in the hypodermis and most other tissues, peaking during the molt (Fig. 3). *ztf-16b::gfp* was expressed in the same cell types and with the same timing during continuous development as during post-dauer development (Fig. 3).

**Figure 3.**
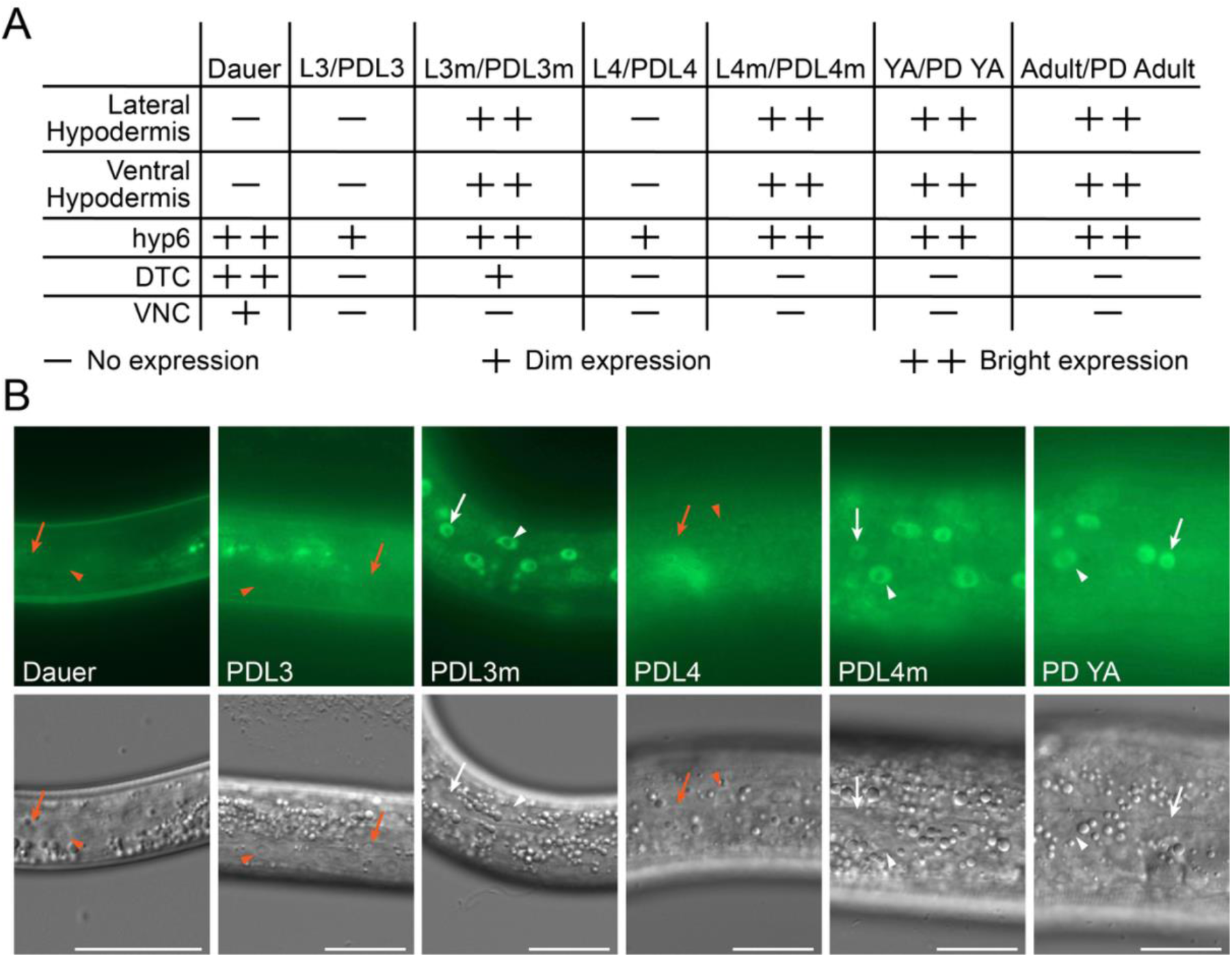
*ztf-16b::gfp* expression oscillates with the molting cycle. (A) *ztf-16b::gfp* is expressed cyclically during development. In the hypodermis, *ztf-16b::gfp* expression increases during the molts and is reduced or absent during intermolt periods. (n = 8-23 worms/stage/life history). Expression of *ztf-16b::gfp* was consistent between individuals at the same timepoint. (B) Representative fluorescent (top) and DIC (bottom) images of the lateral hypodermis. Only autofluorescence was observed in the lateral and ventral hypodermis in dauer, post-dauer L3 (PDL3), and PDL4 stages, while nuclear *ztf-16b::gfp* expression was seen during the PDL3 molt (PDL3m), PDL4m, and post-dauer young adult (PD YA) stages. Representative nuclei lacking *ztf-16b::gfp* expression are indicated by orange arrows (seam cells) and arrowheads (hyp7 nuclei). White arrows/arrowheads indicate the same cell types expressing *ztf-16b::gfp*. In all cases, the scale bar = 20 μm. Images of dauer larvae were taken with a 63X objective whereas all other stages were taken with a 40X objective. “Young adults” did not have embryos whereas “adults” were gravid; DTC, distal tip cell; lateral hypodermis, seam cells and hyp7 nuclei; ventral hypodermis, P cell descendants that have differentiated and fused with the surrounding hyp7; VNC, ventral nerve cord.

In addition to the endogenous *ztf-16b::gfp*, we examined the expression from the transgene present in the rescuing fosmid, *wgIs494[ztf-16b::egfp],* where *egfp* is fused to the C-terminal end of the *ztf-16b* isoform. This transgene was expressed cyclically in the hypodermis, similar to endogenous *ztf-16b::gfp* (Fig. S2). The cyclical expression of *ztf-16b::gfp* appears to be regulated at least partially at the transcriptional level because in both life histories, a transcriptional reporter of *ztf-16* showed increased seam cell expression during the L3 molt compared to the L3 and L4 intermolt periods (Table S5). This finding is consistent with the oscillation found in *ztf-16* mRNA levels during continuous development (Hendriks et al. 2014). ***ztf-16* acts downstream of *let-7***

The major phenotype observed in *ztf-16* mutants is the precocious expression of *col-19p::gfp*. During continuous development, *col-19p::gfp* expression is directly regulated by LIN-29, which is in turn indirectly regulated by *let-7* (Liu et al. 1995; Rougvie and Ambros 1995; Reinhart et al. 2000; Slack et al. 2000; Aeschimann et al. 2019). To determine if *ztf-16* interacts with these genes, we performed a series of epistatic experiments. We first examined the impact of the temperature sensitive *let-7(n2853)* allele on the precocious expression of *col-19p::gfp* in *ztf-16(tm2127)* mutants. *col-19p::gfp* expression was categorized as “Bright,” “Moderate,” or “Dim” when expressed or “Off” when not expressed (Fig. S1). These categories were used for all *col-19p::gfp* expression experiments. When raised at restrictive temperatures, we found that the *let-7* mutation had a minimal impact on precocious *col-19p::gfp* expression in *ztf-16* mutants (Fig. 4A). At the PDL4 stage, *ztf-16 let-7* double mutants retained precocious *col-19p::gfp* expression, albeit at slightly dimmer levels when compared to *ztf-16* mutants (Fig. 4A, left). During continuous development, the loss of *let-7* had no obvious effect as *ztf-16 let-7* double mutants displayed strong, precocious *col-19p::gfp* expression at levels comparable to *ztf-16* single mutants (Fig. 4A, right). With the caveat that *let-7(n2853)* is not a null allele (Reinhart et al. 2000), our data suggest that in larvae, *ztf-16* blocks *col-19p::gfp* expression largely independently of *let-7*.

**Figure 4.**
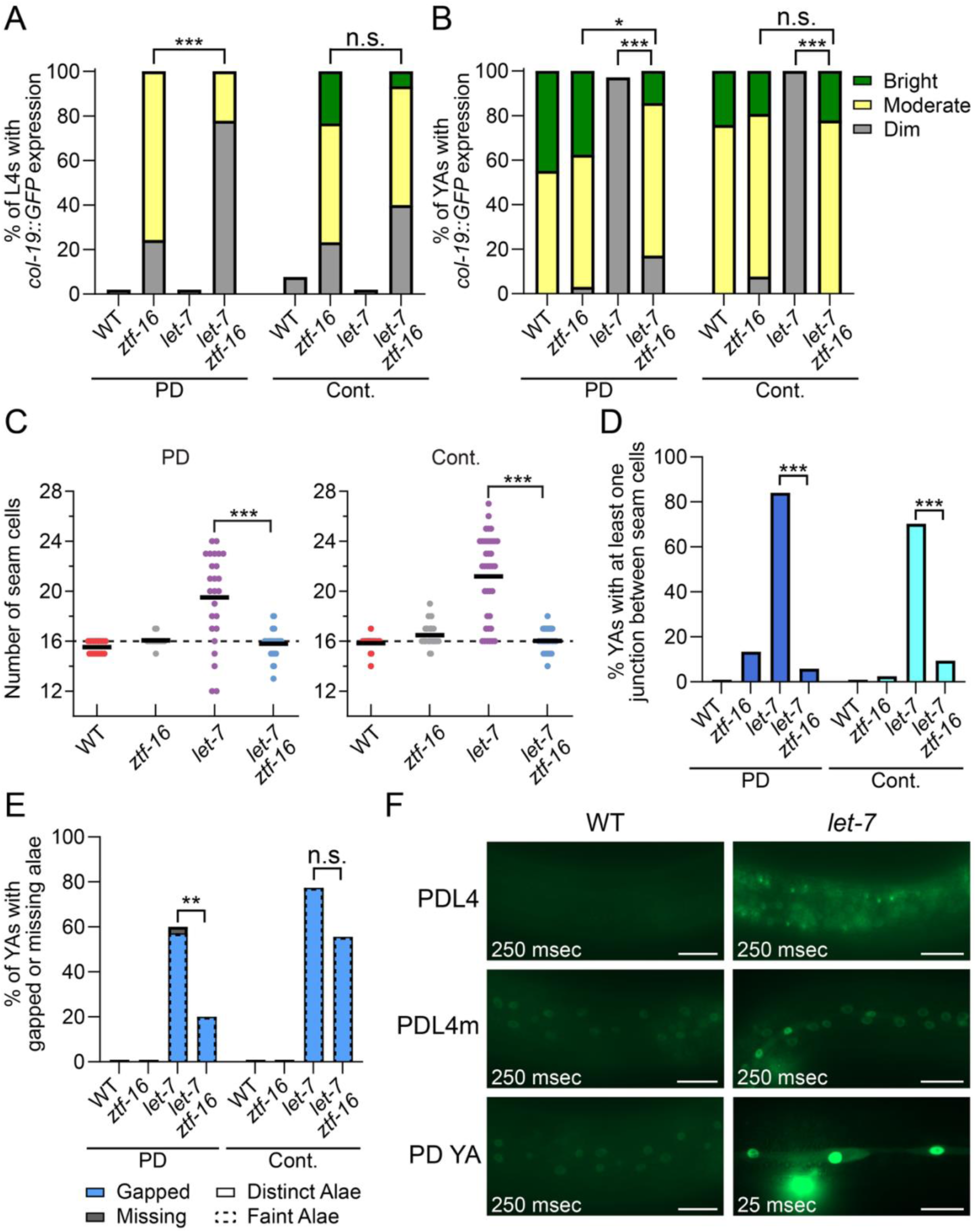
*ztf-16(-)* mutations suppress *let-7(n2853)* phenotypes to varying extents. (A and B*) ztf-16 let-7* double mutants and *ztf-16* single mutants displayed similar levels of *col-19p::gfp* expression during L4 (A) and young adult (B) stages (n = 26-37 worms/strain/stage/life history). (A) *let-7* modestly suppressed precocious *col-19p::gfp* expression in PDL4 and had no effect in continuous L4. Exposure =30 msec. (B) In *let-7* mutant adults, *col-19p::gfp* expression was present but dim in seam cells and absent from hyp7 nuclei (Exposure = 10 msec). Expression was largely restored in *ztf-16 let-7* double mutants. (C) The extra seam cell phenotype observed in *let-7* mutants was suppressed by *ztf-16.* Seam cells were counted using *wIs78[scm::gfp, ajm-1::gfp]* (n = 15-39 worms/strain/life history). (D) The *let-7* seam cell fusion phenotype was suppressed by *ztf-16.* Worms were scored positive for junctions if at least one junction formed between adjacent seam cells. *let-7* mutant worms had 1-9 junctions. (n = 15-40 worms/strain/life history). (E) The *let-7* gapped alae phenotype was moderately suppressed in the *ztf-16 let-7* double mutant. However, the poor quality of alae in the *let-7* mutant was essentially unaffected by *ztf-16.* (F) *ztf-16b::gfp* was overexpressed in *let-7(n2853)* mutants compared to wild type, particularly in the seam cells. (n = 12-20 worms/stage/life history). Scale bar = 20 μm. *ztf-16*, *ztf-16(tm2127); let-7*, *let-7(n2853)* for all genotypes. In panels A, B, and E all strains also contain *maIs105[col-19p::gfp]*. *** p < 0.001, ** p < 0.01, * p < 0.05, and n.s. (not significant, p > 0.05). A two-tailed *t* test was used for panel C; significance was assessed using the Fisher Exact Test for all other panels.

We next asked how the loss of *ztf-16* impacts the reiterative *let-7(-)* phenotypes seen in adults. Prior to answering this question, we needed to establish which, if any, *let-7(n2853)* reiterative phenotypes are present after dauer, since the phenotypes of many heterochronic genes are suppressed after dauer (Liu and Ambros 1991; Abrahante et al. 1998; Abrahante et al. 2003; Karp and Ambros 2012). However, most of the genes that show post-dauer suppression act relatively early in development, prior to dauer formation. Reiterative phenotypes caused by loss of the late-acting heterochronic gene *lin-29* are not suppressed post-dauer (Liu and Ambros 1991), raising the possibility that *let-7(-)* worms may also display reiterative phenotypes after dauer. We found that all of the *let-7(n2853)* reiterative phenotypes tested were also observed post-dauer (Fig. 4). As described below, the loss of *ztf-16* suppressed each of these phenotypes to varying extents in both life histories.

In young adult *let-7(n2853)* mutants raised at restrictive temperatures, *col-19p::gfp* expression is rare in hyp7 nuclei and present but greatly diminished in seam cells (Hurschler et al. 2011; Chu et al. 2014; Perales et al. 2014) (Fig. 4B). These *col-19p::gfp* phenotypes were also observed after dauer (Fig. 4B, left). By contrast, *ztf-16* mutants expressed *col-19p::gfp* at wild-type levels at the young adult stage (Fig. 4B). Loss of *ztf-16* suppressed the *let-7* phenotype; in the double mutant, *col-19p::gfp* was restored in seam cells and hyp7 in both life histories, although expression was slightly dimmer after dauer (Fig. 4B).

As *let-7* affects many aspects of adult cell fate, we next asked if the loss of *ztf-16* could suppress the other reiterative *let-7(-)* phenotypes. Wild type and *ztf-16* mutant adults have 16 seam cells regardless of life history (Sulston and Horvitz 1977) (Fig. 4C). On the other hand, *let-7* mutants undergo an extra seam cell division after the L4 molt (Reinhart et al. 2000). This division can cause seam cell number to increase to 20 or more (Hayes et al. 2006; Chan and Slack 2009; Aeschimann et al. 2019). We observed that more than three-quarters of young adult *let-7* mutants had more than 16 seam cells and that this phenotype was similar in both life histories. In both cases, this phenotype was efficiently suppressed by loss of *ztf-16* (Fig. 4C).

Consistent with the increase in seam cell number, we saw evidence of ongoing seam cell division in *let-7(-)* adults. In wild-type larvae, seam cells are connected to each other and to the surrounding hyp7 syncytium by apical junctions which can be visualized with the apical junction marker *ajm-1::gfp* (Podbilewicz and White 1994; Mohler et al. 1998; Koh and Rothman 2001). By the L4m, seam cells exit the cell division cycle and fuse with each other, while apical junctions are still present between the seam cell syncytium and hyp7 (Podbilewicz and White 1994). When these adult seam cells divide in *let-7* mutants, *ajm-1::gfp* is visible at a point between the two newly forming daughter cells. We observed this focal *ajm-1::gfp* expression in nearly two thirds of *let-7(-)* young adults (Fig. S7, 4D). However, this phenotype was largely suppressed in *ztf-16 let-7* double mutants (Fig. 4D).

During these experiments, we noticed that *ztf-16* mutants displayed an additional seam cell phenotype where a small break was present between the *ajm-1::gfp*-labeled junctions between adjacent seam cells (Fig. S8). These breaks appeared smaller than similar breaks that were previously described in mutants with defects in hypodermal morphogenesis or seam cell fate (Fujii et al. 2002; Smith et al. 2005; Brabin et al. 2011). The basis for this phenotype is unclear and was not explored further.

Adult alae are a cuticular structure secreted from adult seam cells and are commonly used as an indicator of adult cell fate (Ambros and Horvitz 1984). *let-7* mutants display alae defects that have been variably described in the literature. The *let-7(mn112)* null allele has been described to either not form alae at all or to have faint and gapped alae at the L4m (Reinhart et al. 2000; Abrahante et al. 2003; Tennessen et al. 2006). In our experience, *let-7(n2853)* mutants raised at restrictive temperatures formed indistinct, gapped alae at the L4m that were difficult to detect but not indistinguishable from the surrounding cuticle (Fig. 4E). The poor quality of alae created by *let-7* mutants may account for some of the discrepancies in the literature regarding its formation at the L4m. As previously mentioned, *ztf-16* mutants do not form precocious alae, and adult alae are distinct and complete, aside from small gaps overlying the breaks between the seam cells. In contrast to the strong suppression of other phenotypes, the loss of *ztf-16* only weakly suppressed the indistinct and gapped alae of *let-7* mutants. Overall, fewer worms contained gaps in their alae (Fig. 4E), suggesting fewer seam cells were displaying a larval cell program. However, this difference was significant only in post-dauer animals (Fig. 4E). This weak suppression suggests that *ztf-16* affects adult alae formation only moderately.

Finally, *let-7(n2853)* mutants grown at restrictive temperatures burst through the vulva after the L4m, resulting in lethality (Reinhart et al. 2000). While *lin-41* is the key target responsible for the bursting phenotype (Slack et al. 2000; Ecsedi et al. 2015), the reduction of a number of transcription factors is also sufficient to reduce or eliminate bursting (Abrahante et al. 2003; Lin et al. 2003; Großhans et al. 2005; Hunter et al. 2013). We found that *ztf-16 let-7* double mutants were viable at adulthood and did not burst, even at restrictive temperatures (Table S6).

### *let-7* regulates *ztf-16b::gfp* expression indirectly

The suppression of *let-7(n2853)* reiterative phenotypes by mutation of *ztf-16* suggests that *ztf-16* acts downstream of *let-7*. Consistent with these findings, *ztf-16* mRNA levels were previously found to be upregulated in *let-7(n2853)* L4 worms during continuous development (Hunter et al. 2013). To test whether *let-7* regulates *ztf-16* expression after dauer, we crossed our endogenous *ztf-16b::gfp* reporter into a *let-7(n2853)* background. In wild-type larvae at PDL4, *ztf-16b::gfp* was undetectable in the hypodermis. By contrast, expression was evident in seam cells and hyp7 nuclei in *let-7* mutants, suggesting a regulatory role for *let-7* (Fig. 4F). Overexpression of *ztf-16b::gfp* was also seen in hypodermal cells near the tail. The extent to which *ztf-16b::gfp* was overexpressed increased at the PDL4m and young adult stages, with the most dramatic overexpression observed in young adults (Fig. 4F). *ztf-16b::gfp* was similarly overexpressed in *let-7(n2853)* mutants that developed continuously (Fig. S9).

Since *let-7* negatively regulates *ztf-16* expression, we wondered whether *let-7* might directly target the *ztf-16* mRNA for silencing. TargetScan (http://www.targetscan.org/worm_52/) predicts a single *let-7* complementary site within the *ztf-16* 3’UTR (Fig. 5A). We used CRISPR Cas9 to alter the seed region of this site within the context of *ztf-16(cen10[ztf-16b::gfp]),* creating two independent alleles with the identical mutation. If this site mediates silencing, then mutation of the site should cause overexpression of *ztf-16b::gfp,* similar to the overexpression observed in *let-7* mutants. However, we found that mutation of the predicted *let-7* complementary site caused no increase in the levels of *ztf-16::gfp* expression in either allele (Figs. 5B, S9A). Taken together, we conclude that *ztf-16* is regulated by *let-7* during both continuous and post-dauer development, and that this regulation is not dependent on the LCS and is likely indirect.

**Figure 5.**
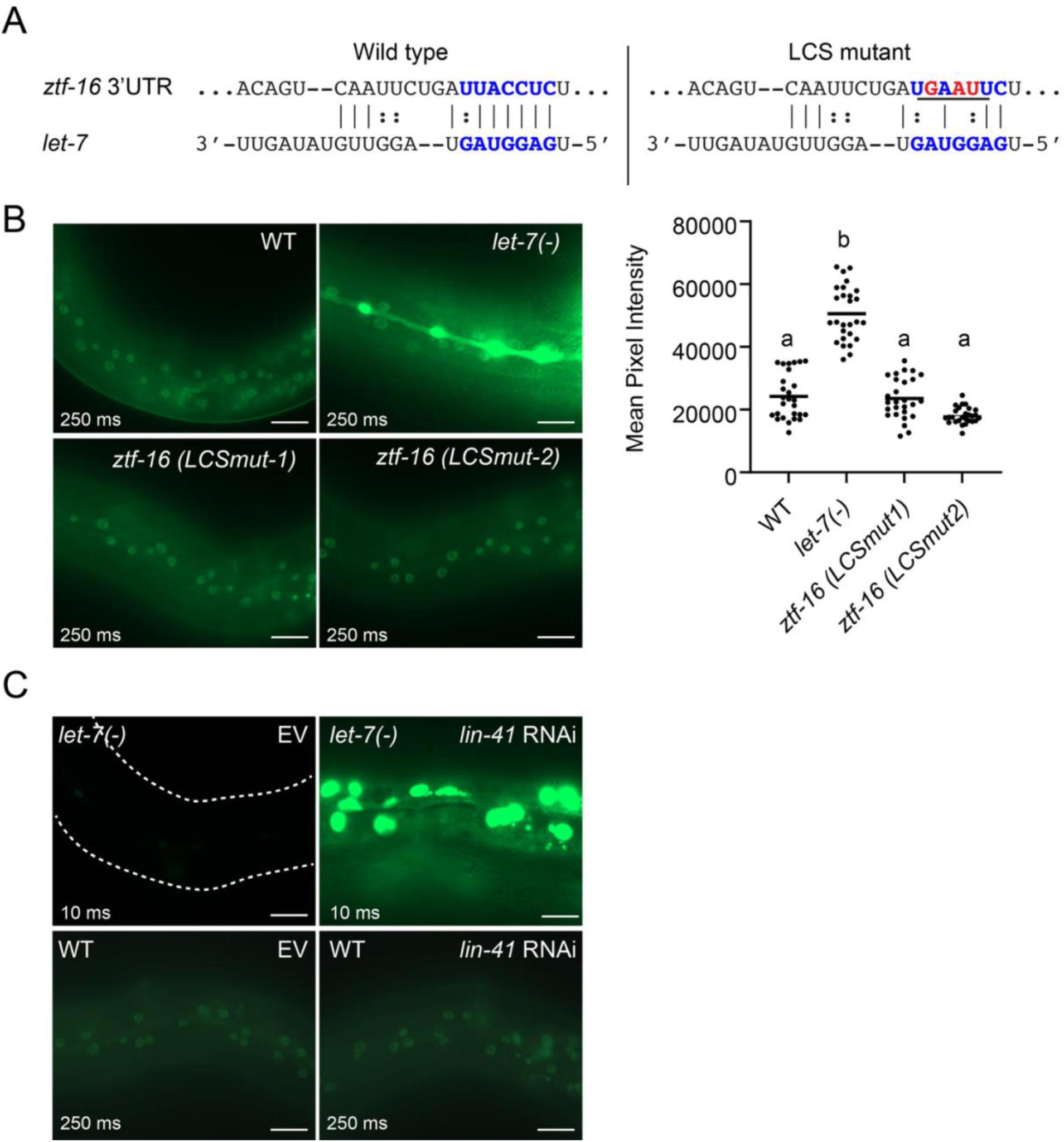
*let-7* regulates *ztf-16* expression indirectly after dauer. (A) A single *let-7* complementary site (LCS) was predicted by TargetScan (http://www.targetscan.org/worm_52/) within the *ztf-16* 3’UTR. The binding of *let-7* to this site was predicted using rna22 (https://cm.jefferson.edu/rna22/Precomputed/GetInputs.jsp) (Miranda et al., 2006). Seed sequence binding is shown in blue bolded letters. Vertical lines represent predicted Watson-Crick base pairing between *let-7* and the *ztf-16* 3’UTR. Dots represent G-U wobble base pairing. We mutated three nucleotides (red) predicted to be bound by the *let-7* seed sequence to create an *EcoRI* site in the genomic DNA (underlined). This *let-7* complementary site mutation (LCSmut) should reduce or eliminate the ability of *let-7* to bind the 3’UTR. (B) We created two independent alleles with the same LCS mutation. Neither allele caused an increase in *ztf-16b::gfp* expression. Representative PDYAs are shown. Image J was used to quantify expression. n = 27 worms/strain. Samples with the same letter indicates no statistical difference between them, and samples with a different letter indicate a statistical difference where signficance = p < 0.05 by Kruskall Wallace and Dunn’s test. Similar results were seen during continuous development (Fig. S9). (C) *let-7* does not downregulate *ztf-16* through regulation of *lin-41.* In fact, *ztf-16::gfp* is more strongly misexpressed in *let-7(-)* PDYAs subject to *lin-41* RNAi (top), while *lin-41* RNAi in a wild-type background does not cause such misexpression (bottom). EV = RNAi treatment with empty vector. n = 20 worms/strain. Scale bars = 20 µm. Exposure times for fluorescent images are listed.

During continuous development, *let-7* mediates its effects on adult cell fate largely via direct silencing of *lin-41,* which encodes an RNA-binding protein that opposes adult cell fate (Reinhart et al. 2000; Slack et al. 2000; Aeschimann et al. 2019). Therefore, one possibility is that *let-7* negatively regulates *lin-41* which in turn positively regulates *ztf-16.* If this were the case, then depletion of *lin-41* should suppress the overexpression of *ztf-16b::gfp* in *let-7* mutants. To test this prediction, we used RNAi to reduce *lin-41* levels in young adult *let-7(n2853)* mutants. Surprisingly, we found that instead of suppression, the overexpression of *ztf-16b::gfp* in *let-7* mutants was strongly enhanced by *lin-41* RNAi (Figs. 5C, S9B). This enhancement was only in the context of *ztf-16b::gfp* expression because, *lin-41* RNAi did suppress the bursting phenotype observed in the *let-7* mutants, consistent with prior work (Slack et al. 2000). In the same experiments where we observed increased *ztf-16::gfp* expression, 94% (208/220) *let-7* mutants treated with empty vector burst whereas only 3% (9/300) *let-7* mutants treated with *lin-41* RNAi burst, demonstrating that the RNAi was functioning correctly. Finally, this effect on *ztf-16b::gfp* expression was only seen in *let-7(-)* mutants; *lin-41* RNAi did not affect *ztf-16b::gfp* levels in wild-type (*let-7(+))* animals (Figs. 5C, S9B). Therefore, the regulation of *ztf-16* by *let-7* occurs neither via direct silencing nor via *lin-41*.

### *ztf-16* appears to act in parallel to *lin-29*

Canonically, the most downstream regulator of adult cell fate is *lin-29.* Other heterochronic genes that produce a precocious phenotype when mutated act directly or indirectly through *lin-29* to regulate the timing of adult cell fate (Ambros 1989; Abrahante et al. 1998; Abrahante et al. 2003; Lin et al. 2003; Aeschimann et al. 2017; Azzi et al. 2020). LIN-29 binds the *col-19* promoter to activate *col-19* expression and acts most directly to promote other adult characteristics such as seam cell fusion, adult alae secretion, and cessation of seam cell divisions (Liu et al. 1995; Rougvie and Ambros 1995). The *lin-29(0)* reiterative phenotypes are not suppressed post-dauer (Liu and Ambros 1991); therefore, it is likely that *lin-29* fulfills this role during both continuous and post-dauer development. We therefore hypothesized that *ztf-16* acts through *lin-29* to regulate *col-19p::gfp* expression. If this hypothesis were correct, loss of *lin-29* should suppress the precocious *col-19p::gfp* phenotype observed in *ztf-16* mutants. Surprisingly, we found that *lin-29(0); ztf-16* double mutants expressed precocious *col-19p::gfp* in both life histories (Fig. 6A). Although the loss of *lin-29* slightly reduced *col-19p::gfp* expression in PDL4 larvae, *col-19p::gfp* was still penetrantly misexpressed in *ztf-16(-)* mutant larvae that lacked *lin-29* (Fig. 6A).

**Figure 6.**
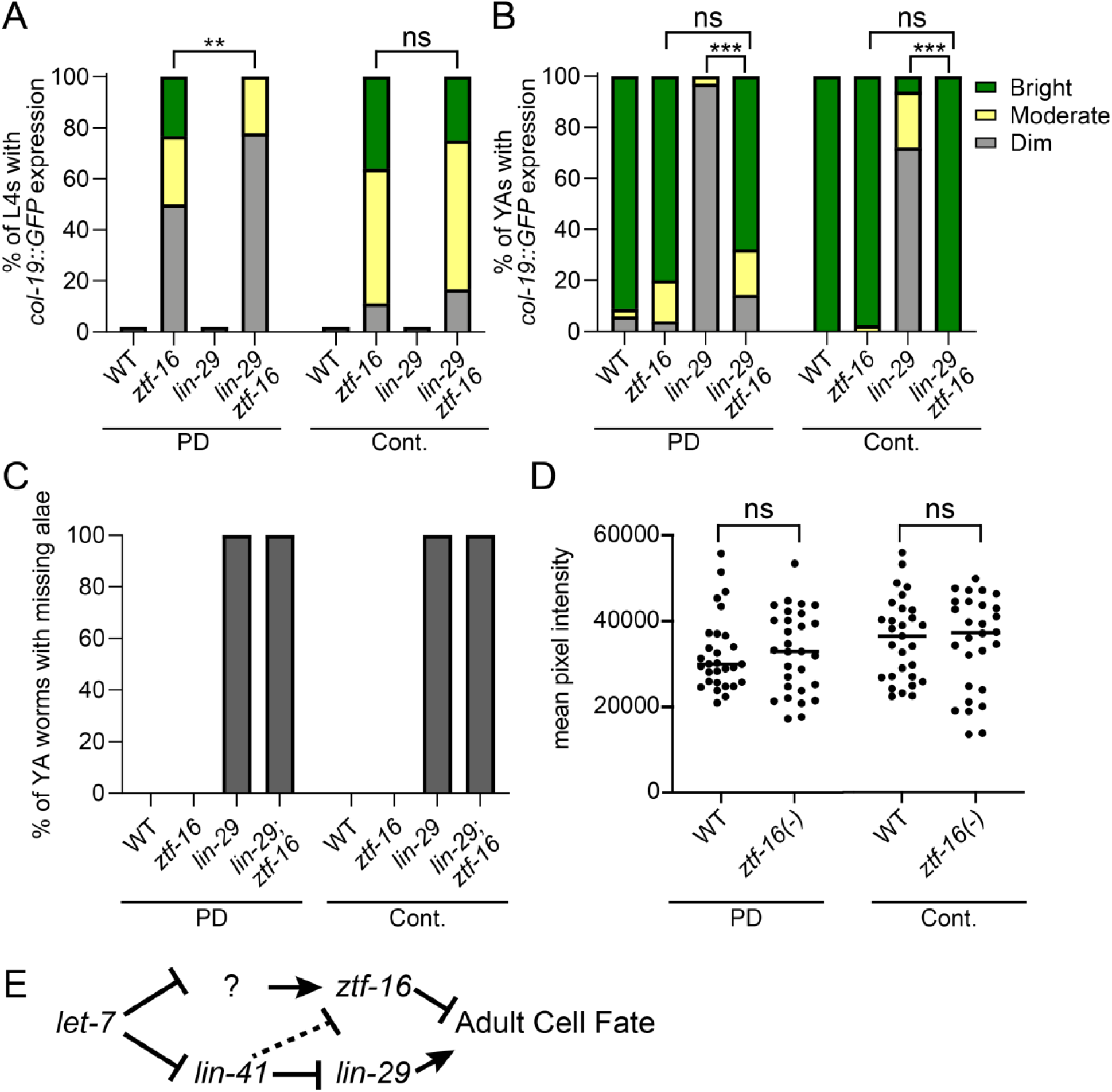
*ztf-16* acts in parallel to *lin-29*. (A) *lin-29(0)* weakly suppresses *col-19p::gfp* expression in *ztf-16* PDL4 larvae and does not suppress *col-19p::gfp* expression in *ztf-16* L4 larvae. (B) *ztf-16* mutant young adults express *col-19p::gfp* even in the absence of *lin-29* (n = 14-50). (C) Mutation of *ztf-16* does not suppress the *lin-29* adult alae phenotype. Alae were either distinct and complete or entirely absent (n = 9-35). (D) Mutation of *ztf-16* does not affect expression of *lin-29(xe61[lin-29::gfp]*. Each dot represents an individual larva at the late PDL3 or continuous L3 stage, when *lin-29::gfp* first becomes visible in seam cells in wild-type and *ztf-16(tm2127)* mutants. (E) Current genetic model of *ztf-16* activity (see text). Statistical significance was assessed using a Fisher Exact Test (A-B) or a Mann Whitney test (D). *** p < 0.001, ** p < 0.01, and ns (not significant), p > 0.05.

We next wondered if the loss of *ztf-16* would suppress the reiterative *lin-29* phenotypes seen in adults. Consistent with the role of LIN-29 as a direct activator of *col-19* transcription, *lin-29(0)* mutants display very dim *col-19p::gfp* expression in continuous and post-dauer adults (Fig. 6B). The residual *col-19p::gfp* expression observed in *lin-29(0)* adults indicates that one or more factors in addition to *lin-29* promotes *col-19p::gfp* expression. We found that *lin-29(0); ztf-16* double mutant adults displayed bright *col-19p::gfp* expression. Therefore, *lin-29* is not required for *col-19p::gfp* expression in *ztf-16* mutants (Fig. 6B)

In addition to regulating *col-19p::gfp, lin-29* is necessary but not sufficient for adult alae formation (Ambros and Horvitz 1984; Abete-Luzi et al. 2020; Azzi et al. 2020). We asked if the loss of *ztf-16* would suppress the lack of adult alae formation in *lin-29* mutants. We found that *lin-29; ztf-16* double mutants did not form adult alae (Fig. 6C). Therefore, *lin-29*, although dispensable for *col-19p::gfp* expression, is still required for adult alae formation in *ztf-16* mutants.

To help clarify the complex relationship between *ztf-16* and *lin-29* implied by the above genetic data, we asked if *ztf-16* can regulate *lin-29* expression. Specifically, we asked if *lin-29* is expressed precociously or at higher levels in *ztf-16* mutants. An endogenous *lin-29::gfp* allele (*xe61)* is brightly expressed in seam cell nuclei during the L4 stage (Aeschimann et al. 2017). If *ztf-16* regulates *lin-29* expression, the loss of *ztf-16* should cause precocious expression of *lin-29::gfp*. To determine if *lin-29::gfp* is expressed precociously in *ztf-16* mutants we examined L3-staged larvae in both life histories. In wild-type animals*, lin-29::gfp* expression in hypodermal cells is first detectable in seam cells in the late L3. Similarly, in *ztf-16* mutants, we did not detect *lin-29::gfp* expression before this stage. At late L3, the levels of GFP expression observed were similar in wild-type and *ztf-16* mutant worms (Fig. 6D). Taken together, these data suggest that *ztf-16* regulates *col-19p::gfp* largely independently of *lin-29*.

## Discussion

We have identified a new heterochronic gene, *ztf-16.* We found that *ztf-16* blocks adult cell fate during both continuous and post-dauer development, primarily by regulating *col-19*, an adult-specific collagen and well-characterized adult cell fate marker. *ztf-16* is the only gene known to oppose adult cell fate equally strongly in both continuous and dauer life histories. In contrast, *lin-14, lin-28, hbl-1, lin-41,* and *lin-42* mutants display strong precocious phenotypes during continuous development, but these phenotypes are partially (*lin-41)* or completely (*lin-14, lin-28, hbl-1, lin-42)* suppressed after dauer (Liu 1990; Liu and Ambros 1991; Abrahante et al. 1998; Abrahante et al. 2003; Spike et al. 2014). Although necessary to block *col-19p::gfp* expression in larvae, *ztf-16* does not appear to be sufficient, because adult worms express both *ztf-16b::gfp* and *col-19p::gfp*.

Most heterochronic genes that oppose adult cell fate are expressed early in development and are subsequently downregulated to allow for later seam cell programs (Rougvie and Moss 2013). By contrast, *ztf-16b::gfp* is expressed cyclically throughout development, with expression peaking during the molts (Fig. 3). Expression does not differ between the dauer and continuous life histories, consistent with the requirement to prevent *col-19p::gfp* expression in both contexts. The cyclical expression pattern of *ztf-16* is similar to the expression of *lin-42*, encoding the ortholog to the Period protein in *Drosophila* and mammals (Jeon et al. 1999). However, the temporal expression pattern is the opposite as *lin-42* levels peak during the intermolt periods and diminish during the molts (Tennessen et al. 2006). *lin-42* regulates the molting cycle and also negatively regulates heterochronic microRNA transcription levels (Monsalve et al. 2011; Perales et al. 2014; Wynsberghe et al. 2014; Edelman et al. 2016). In contrast, no molting defects have been observed in *ztf-16* mutants, suggesting that *ztf-16* does not regulate the molting process, despite cyclical expression.

*ztf-16* interacts with the heterochronic pathway in a novel manner. Canonically, adult cell fate, including *col-19p::gfp* expression, is most directly regulated by the *lin-29* transcription factor (Ambros 1989; Liu et al. 1995; Rougvie and Ambros 1995). Early in development, translation of the *lin-29a* isoform is repressed by the LIN-41 RNA-binding protein (Slack et al. 2000; Aeschimann et al. 2017; Azzi et al. 2020). Later in development, the *let-7* microRNA directly silences *lin-41,* thereby allowing *lin-29* expression (Reinhart et al. 2000; Aeschimann et al. 2019). Our experiments suggest that *ztf-16* acts downstream of *let-7,* but in parallel to *lin-29* (Fig. 6E). First, genetic epistasis experiments showed that *ztf-16* suppressed all of the *let-7* reiterative defects, to varying extents. Furthermore, endogenous *ztf-16b::gfp* was strongly misexpressed in a *let-7(-)* background. This regulation was not mediated via the single predicted *let-7* complementary site in the *ztf-16* 3’UTR, suggesting that *let-7* regulates *ztf-16* indirectly. The *let-7-*mediated downregulation of *ztf-16* also did not occur via *lin-41,* and in fact, *lin-41* antagonizes *ztf-16* expression because *ztf-16b::gfp* was more strongly misexpressed when both *let-7* and *lin-41* were depleted. This antagonistic relationship is surprising because *lin-41* and *ztf-16* act in the same direction within the heterochronic pathway: both genes oppose adult cell fate. Furthermore, *lin-41(-)* precocious phenotypes present during continuous development are partially suppressed after dauer (Liu 1990; Spike et al. 2014); however, ZTF-16b::GFP expression was strongly affected by *lin-41* RNAi in both continuous and dauer life histories (Figs. 5, S9). Finally, we showed that *lin-29* was dispensable for *col-19p::gfp* expression in *ztf-16(-)* larvae and adults, and that *ztf-16* did not affect expression of endogenous *lin-29::gfp*.

Since LIN-29 directly activates expression of the *col-19* promoter (Liu et al. 1995; Rougvie and Ambros 1995), we hypothesize that *ztf-16* acts in parallel to *lin-29* to regulate this aspect of adult cell fate (Fig. 6E). One possibility is that ZTF-16 may bind to the *col-19* promoter and directly repress *col-19* expression. Alternatively, *ztf-16* may regulate *col-19* expression indirectly.

The NHR-25 nuclear hormone receptor has also been implicated in *lin-29-*independent regulation of *col-19p::gfp* expression*. nhr-25(-)* L4 staged larvae exhibit strong *col-19p::gfp* expression, even in the presence of a *lin-29* null mutation (Hada et al. 2010). Surprisingly, despite this precocious *col-19p::gfp* phenotype, RNAi of *nhr-25* suppresses the precocious formation of adult alae of various heterochronic mutants, suggesting *nhr-25* has opposing regulatory roles on *col-19p::gfp* and adult alae (Hada et al. 2010). *ztf-16* also affects alae and *col-19p::gfp* differently, but for *ztf-16* the difference is in degree rather than direction. *ztf-16* appears to have a larger role in regulating *col-19p::gfp* than in regulating alae. *ztf-16* mutants express *col-19p::gfp* strongly in larvae and adults, even in *let-7* and *lin-29* mutant backgrounds.

By contrast, *ztf-16* mutants do not display precocious alae, show only weak suppression of the gapped alae phenotype of *let-7* mutants, and do not affect alae formation in *lin-29* mutants. Notably, we previously described another example of *lin-29-*independent regulation of *col-19p::gfp* (Wirick et al. 2021). *daf-16* promotes dauer formation (Vowels and Thomas 1992). We found that *daf-16* is required to block *col-19p::gfp* expression in dauer larvae, thereby coordinating the decision to enter dauer with the opposition of adult cell fate. Unlike *ztf-16, daf-16* regulates *col-19p::gfp* expression upstream of *lin-41*. When either *daf-16* or *lin-41* are depleted, dauer larvae express *col-19p::gfp,* even in the presence of a *lin-29* null allele (Wirick et al. 2021). DAF-16 does not bind the *col-19* promoter (Wirick et al. 2021), and *col-19p::gfp* is a transcriptional reporter and is therefore unlikely to be directly regulated by the LIN-41 RNA-binding protein, suggesting that unknown transcription factors act downstream of *daf-16* and *lin-41*. ZTF-16 is unlikely to be the transcription factor in question for two reasons. First, *ztf-16* does not affect *col-19p::gfp* expression during the dauer stage (Fig 1B). Second, RNA-seq data comparing *daf-16(0)* mutant to control dauer larvae found that *ztf-16* expression is not significantly affected (Wirick et al. 2021). It will be interesting to determine whether the unknown transcription factors that regulate *col-19p::gfp* expression in *daf-16* mutant dauer larvae can also regulate *col-19p::gfp* expression in *ztf-16* mutants.

## Supporting information

RNAseq Dataset

## Acknowledgements

We are grateful to Victor Ambros (University of Massachusetts Medical School) in whose lab the original screen was performed. We thank former undergraduate students at Central Michigan University who contributed to backcrossing and other aspects of this work, including Stephen Domingue, Benjamin Prout, Brittany Lardie, and Amberlyn Hales. We thank Jennifer Schisa (Central Michigan University) and lab members for helpful discussions. We thank WormBase for information (Sternberg et al. 2024). Some strains were provided by the Caenorhabditis Genetics Center (CGC), which is funded by NIH Office of Research Infrastructure Programs (P40 OD010440). DNA quality analyses, Illumina library prep, WGS sequencing and primary analysis were conducted at the Genetic Resources Core Facility, Johns Hopkins Institute of Genetic Medicine, Baltimore, MD. This work was supported by NIH R01HD109667 and R35GM148248 (JKK), NIH R01GM050227 and P40OD010440 (AR), NIH R35GM124828 (AZ), NSF CAREER 1652283 and NIH R15GM117568 and R15GM150082 (XK).

