## Supplemental Figures and Tables for "*ztf-16* is a novel heterochronic modulator that opposes adult cell fate in dauer and continuous life histories in *Caenorhabditis elegans*"

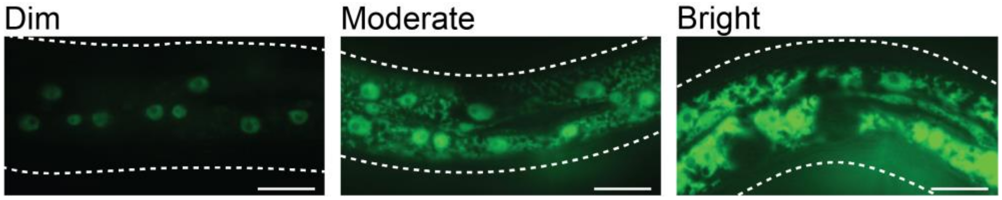


**Figure S1. Categorization of *col-19p::gfp* expression.** Detectable *col-19p::gfp* expression was categorized as dim, moderate, or bright. As explained in the Methods section, three images were captured of each worm and used sort worms into phenotypic categories. The above images show varying *col-19p::gfp* expression in wild-type young adults (Exposure = 10 msec). Note that at this short exposure time, no autofluorescence is observed; all of the fluorescence is GFP. Scale bar = 20 μm.

| | L3 | L3m | L4 | L4m | YA |
|---|---|---|---|---|---|
| Lateral Hypodermis | — | ++ | — | ++ | ++ |
| hyp6 | + | ++ | + | ++ | ++ |

— No expression + Dim expression ++ Bright expression

**Figure S2. *wgIs494[ztf-16b::TY1::egfp]* is expressed cyclically in the hypodermis.** *wgIs494* was created by modENCODE and contains recombineered WRM0633cF04, a 31kb fosmid that includes the *ztf-16* genomic region plus the upstream and downstream genes, *kola-2* and *mig-13,* respectively (Gerstein *et al.* 2010). The *b* isoform of ZTF-16 was tagged with EGFP (Sarov *et al.* 2006). This transgene rescued *ztf-16* mutants (**Figure 1C**). YA, young adult (no embryos).

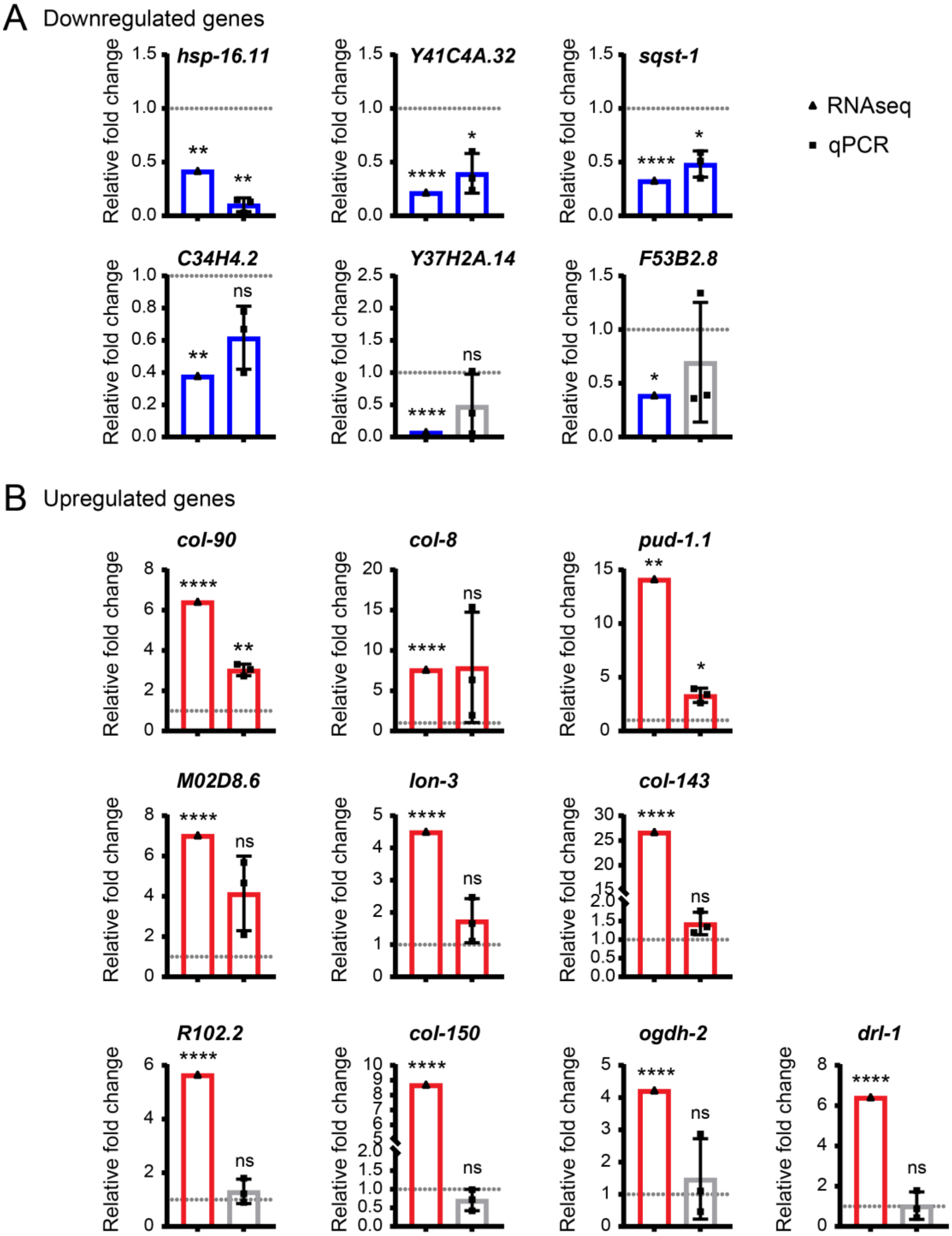


**Figure S3. qPCR validation of RNA-seq data.** A selection of genes that changed substantially (>2x) and significantly (FDR < 0.05) in *ztf-16(tm2127)* mutant PDL4 larvae vs. wild-type control PDL4 larvae based on RNA-seq data (triangles, left) were tested by qPCR (squares, right). For RNA-seq, the fold change was derived from the log2FoldChange in File S1 (FC = 2^(log2FC)). For qPCR, PDL4 larvae were handpicked at the L4.5-L4.7 stage to ensure that differences in gene expression were not due to differences in staging between samples. Three biological replicates from samples grown on different days were collected. Fold change was calculated by the ΔΔCT method, where samples were normalized to the *eft-2* housekeeping gene. Dashed lines indicate FC = 1, or no difference between the samples. Downregulated genes are shown in blue and upregulated genes are shown in red. qPCR data were considered down- or upregulated if all 3 biological replicates changed in the same direction. Statistical significance indicates FDR (RNA-seq data) or was determined by paired t-test (qPCR) data. **** $p < 0.001$, ** $0.001 < p < 0.01$, * $0.01 < p < 0.05$, ns $p \geq 0.05$.

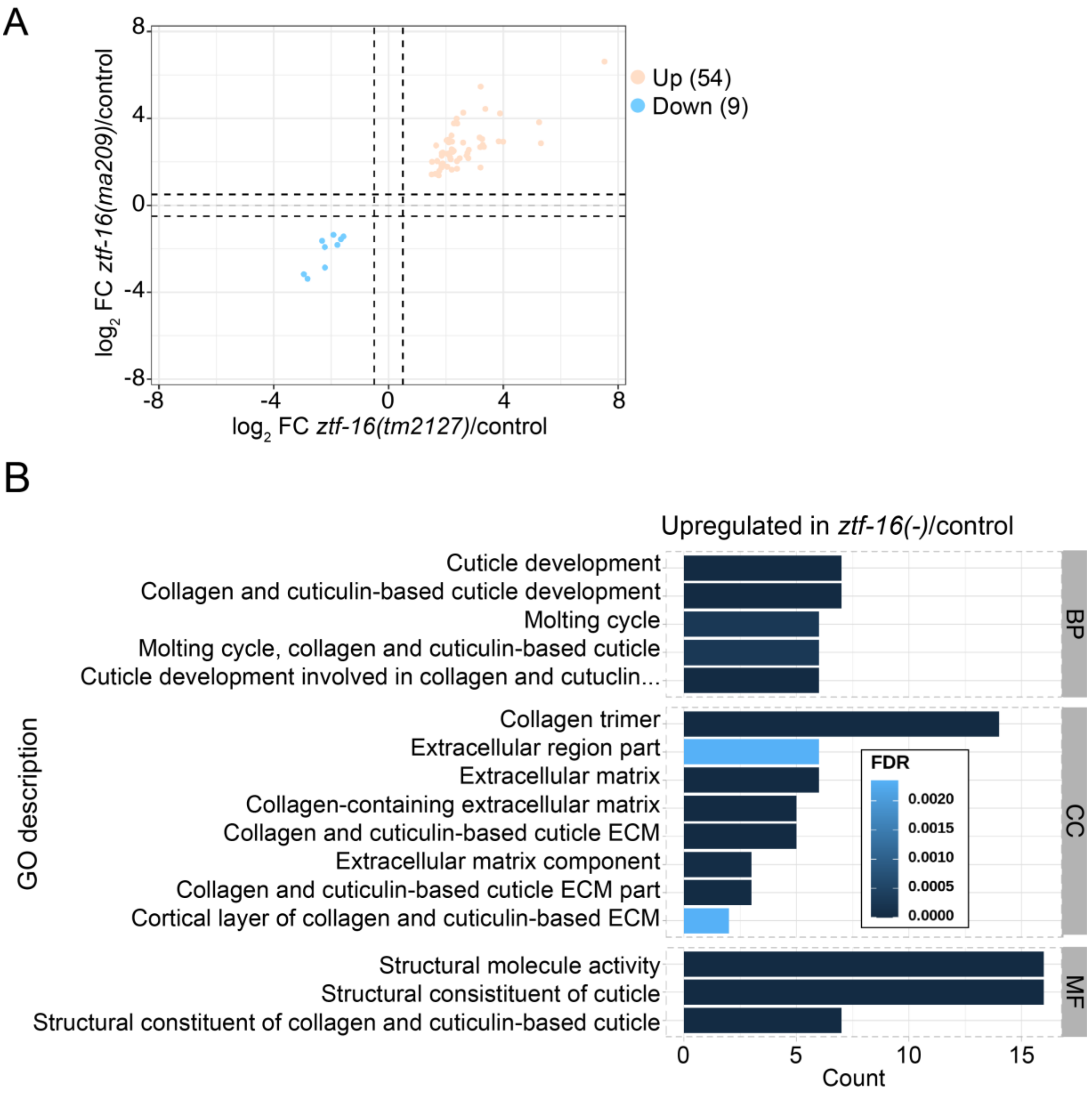


**Figure S4: Gene expression changes in *ztf-16* mutant larvae during continuous development**. mRNA-seq data were collected from two different *ztf-16* mutants and wild-type controls during the L4 stage. (A) A four-factor plot showing all genes whose expression changed significantly (FDR ≤ 0.05) in both mutants relative to controls. Genes with a $\log_2$ fold change (FC) ≥ 0.5 in both mutants were considered upregulated and are shown in orange. Genes with a $\log_2$ fold change ≤ -0.5 in both mutants were considered downregulated and are shown in blue. (B) Significantly (FDR ≤ 0.05) enriched GO terms associated with the upregulated genes. There were too few downregulated genes for associated GO-terms to be significantly enriched; however, the GO-terms ascribed to these genes are listed in File S1, along with a complete list of genes, $\log_2$ fold-change, FDR, and enriched GO terms.

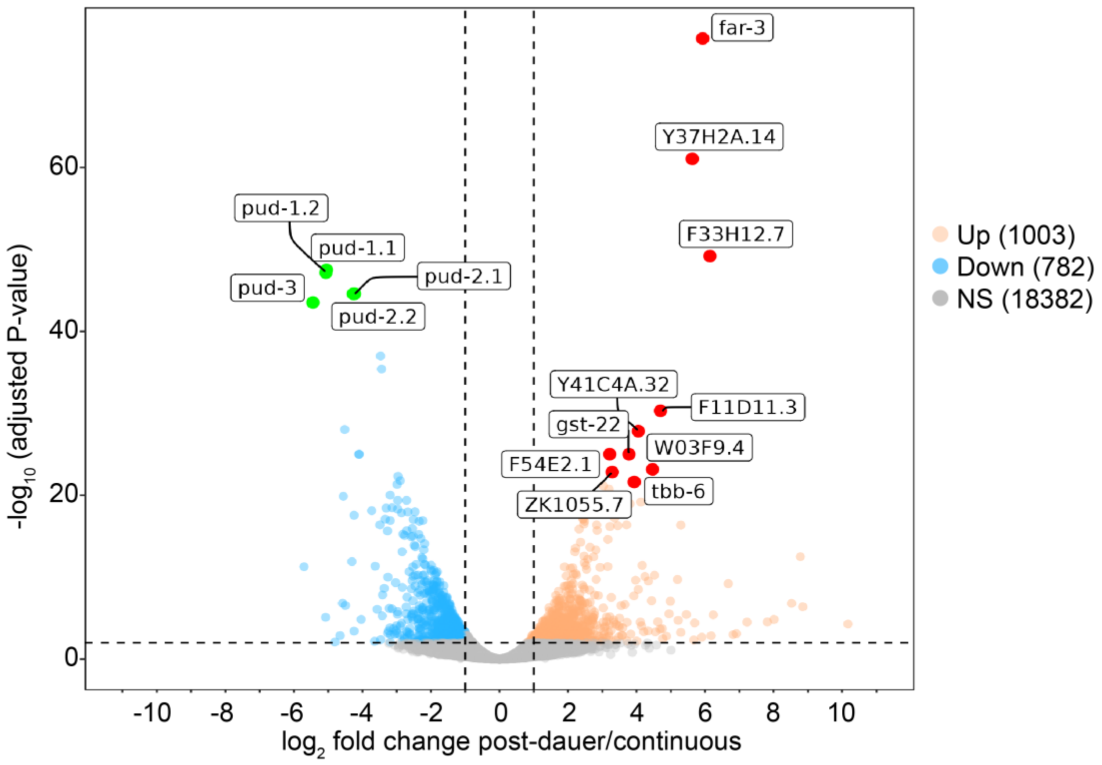


**Figure S5. Gene expression changes in wild-type post-dauer larvae.** Volcano plot depicting DEG analysis of mRNA-seq data from wild-type post-dauer L4 larvae vs. wild-type L4 larvae developing continuously. Genes displaying a false discovery rate (FDR) of ≤ 0.05 and a $\log_2$ fold change ≥1 are shown in orange, whereas genes displaying an FDR of ≤ 0.05 and a $\log_2$ fold change ≤ -1 are shown in blue. All other genes are shown in gray. The most significantly affected genes are further highlighted in red (upregulated genes) or green (downregulated genes).

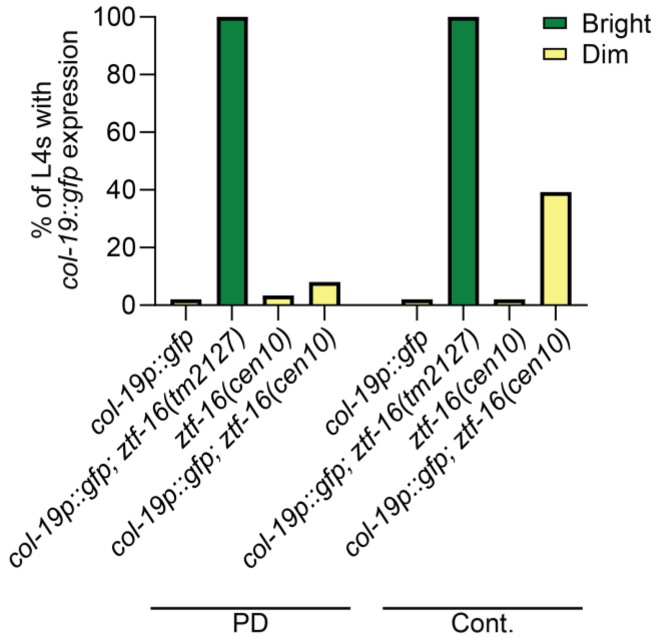


**Figure S6. Endogenously-tagged *ztf-16* is functional.** Precocious expression of *col-19p::gfp* was used to indicate compromised *ztf-16* function. *ztf-16(cen10[ztf-16b::gfp])* larvae rarely displayed precocious *col-19p::gfp*. In the cases when expression was observed, this expression was dim and in only a subset of hypodermal nuclei. By contrast, *ztf-16(tm2127)* mutants display bright *col-19p::gfp* expression throughout the hypodermis and with complete penetrance. *ztf-16b::gfp* and *col-19p::gfp* were distinguished from one another in two ways. First, *ztf-16b::gfp* is not expressed in the lateral hypodermis during the L4 stage (Figure 3). Second, even during stages when *ztf-16b::gfp* expression is visible, an exposure time of 250ms is needed for reliable visualization. Thus, the exposure time of 30ms used in this experiment should be insufficient to view *ztf-16b::gfp*, while still capturing the bright *col-19p::gfp* expression. As a control, larvae containing *ztf-16(cen10)* but lacking *col-19p::gfp* were scored under the same conditions, as shown above. (n = 28-40 worms/strain/life history).

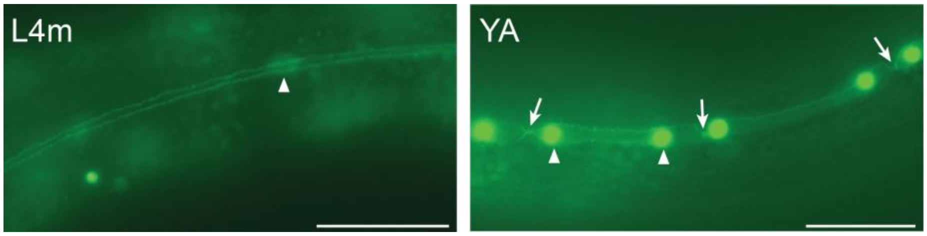


**Figure S7. Junctions re-emerge between seam cells in *let-7* mutant young adults**
In wild-type worms, seam cells begin fusing in the L4 stage and are completely fused by the L4m. In *let-7(n2853)* mutants, seam cells fuse normally at restrictive temperatures at the L4m, as visualized by *wIs78[SCMp::gfp, ajm-1::gfp]* (left). Junctions (indicated by white arrows) re-emerge in young adults (right). White arrowheads indicate seam cell nuclei. The re-emergence of junctions may be explained by an inappropriate division of the seam cells (Azzi *et al.* 2020). Scale bar = 20 µm.

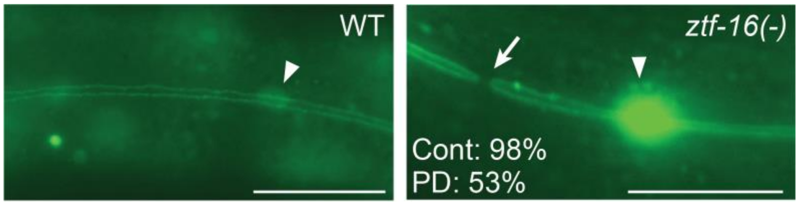


**Figure S8. *ztf-16* mutants have a small break between adjacent seam cells after the L4m.** In wild type, by the L4m, seam cells have fused, forming a syncytium, visualized by *wIs78[SCMp::gfp, ajm-1::gfp]* (left). While scoring seam cell fusion in young adults, we noticed that small breaks occurred between adjacent seam cells in *ztf-16* mutants during both life histories (white arrow, right). Alae were absent from the cuticle overlying the small space, consistent with the role for seam cells in secreting alae. Gaps between seam cells have been observed in situations where epidermal morphogenesis is disrupted (Fujii *et al.* 2002; Smith *et al.* 2005; Brabin *et al.* 2011). In the absence of *plx-1* activity, seam cell boundaries close midway in the seam cell, resulting in a large space between adjacent seam cells and the lack of alae above the gap (Fujii *et al.* 2002). When subjected to *elt-1* RNAi, seam cells lose their identity as seam cells and fuse with the hyp7 in a division-independent manner, resulting in a decrease in seam cell number (Smith *et al.* 2005; Brabin *et al.* 2011). The breaks observed in *ztf-16* mutants appear much smaller than the breaks mentioned above, and *ztf-16* mutant adults have the correct number of seam cells (Figure 4). The basis for this phenotype is unclear and was not explored further (n = 15-40 worms/life history). Percentages indicate the percentage of worms that had a least one break between adjacent seam cells. Individual worms contained 1-6 breaks. White arrowheads indicate seam cell nuclei. Scale bar = 20 μm.

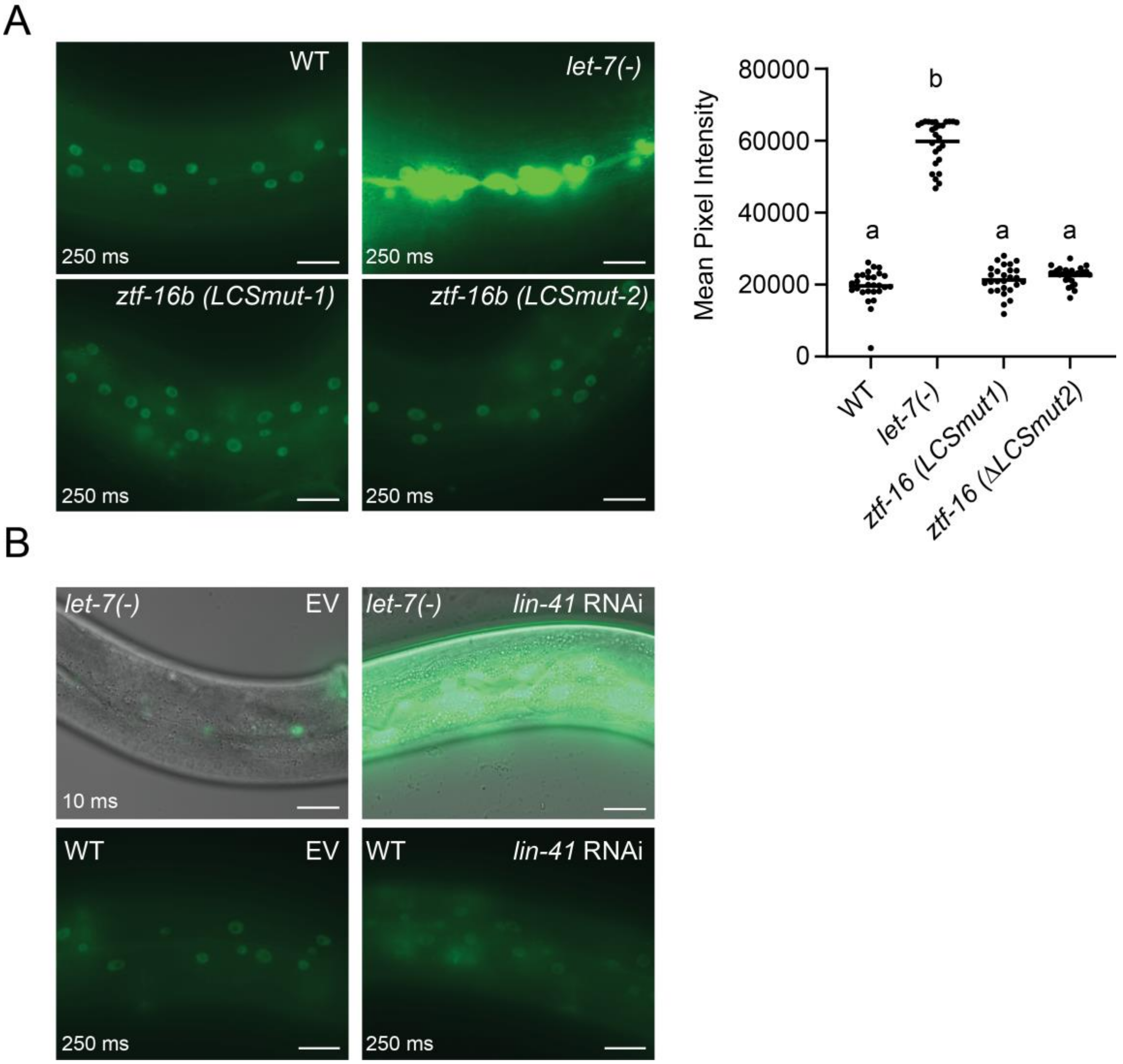


**Figure S9. *let-7* regulates *ztf-16* expression indirectly during continuous development**. (A) We created two independent alleles with the same *let-7* complementary site (LCS) mutation (see Fig. 5A). Neither allele caused an increase in *ztf-16b::gfp* expression. Representative YAs are shown. Image J was used to quantify expression. n = 27 worms/strain. Samples with the same letter indicates no statistical difference between them, and samples with a different letter indicate a statistical difference where signficance = $p < 0.05$ by Kruskall Wallace and Dunn's test. (B) *let-7* does not downregulate *ztf-16* through regulation of *lin-41.* In fact, *ztf-16::gfp* is more strongly misexpressed in *let-7(-)* YAs subject to *lin-41* RNAi (top), while *lin-41* RNAi in a wild-type background does not cause such misexpression (bottom). EV = RNAi treatment with empty vector. n = 20 worms/strain. Scale bars = 20 μm. Exposure times for fluorescent images are listed.

| Allele | Oligo Name | Sequence 5' → 3' |
|---|---|---|
| *cen10*, Forward 1 | XK142 | AGT CAA CAC TCC GTG AAC CTT |
| *cen10*, Reverse 2 | XK143 | CGC CTC CTT CTT CCC TCA ATT T |
| *cen10*, Reverse 1 | XK144 | CTT TTC TCT TCT CCC ACT CGC T |
| *cen10*, Forward 2 | XK145 | CCC TTT CCT TAC GGG AGA TCA G |
| *cen10*, Sequencing | XK150 | CTT TTC TCT TCT CCC ACT CGC T |
| *cen10*, Sequencing | XK151 | CCC TTT CCT TAC GGG AGA TCA G |
| *ma209*, Forward 1 | XK30 | TGT GAT ATC CTG AAC GTG AAC TCT |
| *ma209*, Reverse 1 | XK31 | ACG AGA AAT TGA AAA CCT CCT CA |
| ma209, Forward 2, Nested | XK32 | TCT GAT TTT CAG TCT TCT CCG AA |
| *ma209*, Reverse 2, Nested | XK33 | CCT CCT CAT CAA ACT ATA CAT GGT G |
| *ma213*, Forward 1 | XK24 | AGT CAA CAC TCC GTG AAC CTT |
| *ma213*, Reverse 1 | XK25 | CGC CTC CTT CTT CCC TCA ATT T |
| *ma213*, Forward 2, Nested | XK26 | CTT TTC TCT TCT CCC ACT CGC T |
| *ma213*, Reverse 2, Nested | XK27 | CCC TTT CCT TAC GGG AGA TCA G |
| *n2853*, Forward 1 | oAA182 | CAG TTC TGC GAC TCC GAA T |
| *n2853*, Reverse 1 | oAA181 | ACC TAG CAG CGG TCG TTT T |
| *n2853,* Forward 1 | XK160 | ACC GAT ACA ACA GTT CTC CGA |
| *n2853*, Reverse 1 | XK161 | CCT CTC GCG GGT TTC TGT TC |
| *sa191*, Forward 1 | XK6 | CAA GCT CAT CGC CAT CTT TGC |
| *sa191*, Reverse 1 | XK7 | AAG CAA ACG TGG GCA ACA GG |
| *sa191*, Forward 2, Nested | XK8 | TTT GCC GTA CTC GTC CTC TC |
| *sa191*, Reverse 2, Nested | XK9 | CGG CAG TGC ATT GAT AGG TG |
| *tm2127*, Forward 1 | AD47 | CGC TTG CAC AGA ATG CGG AT |
| *tm2127*, Reverse 1 | AD48 | GGC AGC GGC TTC TTA CGC AA |
| *tm2127*, Forward 2 | AD49 | GCG GAT TCA CGA CTA CTG TA |
| *tm2127*, Reverse 2 mut | AD50 | GCC GTT AAA TGG TGG GGC TA |
| *tm2127*, Reverse 1 WT | XK141 | AGA TAC GGG GGC AGT ACA AA |
| *tm2127,* Forward 3 | XK210 | GGC ACC AAA GGA CAA AAG AC |
| *tm2127,* Reverse 3 | XK211 | ATC TGT TTG CCC AAA ACC GT |

**Table S1. Primers used for genotyping.**

| Oligo Name | Sequence 5’ → 3’ | Role |
|---|---|---|
| XK135 | tcttgCGTTGCTTGAGTACTATGAT | Guide 1, Forward |
| XK136 | aaacATCATAGTACTCAAGCAACGc | Guide 1, Reverse |
| XK137 | tcttgTGGATGATGGTGAATTCAGG | Guide 2, Forward |
| XK138 | aaacCCTGAATTCACCATCATCCAc | Guide 2, Reverse |
| XK106 | ggctgctcttcgtggTGGATTCTCTACATCACC | 5’ Homology Arm, Forward |
| XK107 | gggtgctcttcgcgcGGCGGAAGTCGTTGCTTGAGTACTAT GATCGACAAAAATGACATGAG | 5’ Homology Arm, Reverse |
| XK108 | ggctgctcttcgacgTGAATTCACCATCATCCAAGAAATATC | 3’ Homology Arm, Forward |
| XK109 | gggtgctcttcgtacTTTGATGAGGGGTGGTGGTG | 3’ Homology Arm, Reverse |

**Table S2. DNA primers used to tag endogenous *ztf-16b* with GFP.**
Two guide RNAs to create a double-stranded break were designed based on highest efficiency and fewest off-site targets using http://crispor.tefor.net/ Guide oligos (XK135-138) were annealed in pairs and ligated into pRB1017 digested by BsaI-V2 (Arribere *et al.* 2014). Homology arms of 500 bp or more were created using XK106-109 and then added to the SapTrap reaction (Schwartz and Jorgensen 2016) to create pMLV416. Uppercase letters reflect bases complementary to endogenous sequences while lower case letters reflect overlaps for ligation (XK135-138) or creation of a SapI digestion site (XK106-109).

| | Hours of Development at 24˚C | |
|---|---|---|
| Stage | Continuous | Post-Dauer |
| L1 | 12 | NA |
| L2 | 20 | NA |
| L3 | 32-36 | 12 |
| L3m | 38 | 16 |
| L4 | 40-42 | 20 |
| L4m | 44 | 24 |
| Young Adult | 50 | 28-30 |
| Gravid Adult | 58 | 38 |

**Table S3. Time required for *C. elegans* to develop to specific stages.** For continuous populations, the time indicates the number of hours after egg-laying. For post-dauer populations, the time indicates the number of hours after dauers have been SDS selected and transferred to a plate with a new food source. m, molt; Young adults have not yet produced embryos, whereas gravid adults contain embryos.

| Relevant Genotype | % PDL4 larvae with precocious *col-19p::GFP* expression | n |
|---|---|---|
| *ma209/+* | 0 | 151 |
| *ma213/+* | 0 | 121 |

**Table S4. *ma209* and *ma213* are recessive.** 6X backcrossed *maIs105 [col-19p::gfp] daf-28; ma209* or *maIs105[col-19p::gfp] daf-28; ma213* hermaphrodites were crossed with males of the following genotype: *arIs131[lag-2p::yfp, ceh-22p::gfp]/+; maIs105[col-19p::gfp] daf-28.* Progeny expressing *ceh-22p::gfp* in the pharynx were observed for precocious *col-19p::gfp* in the hypodermis at the PDL4 stage.

| Stage | GFP expression in seam cells | GFP expression in hyp7 nuclei |
|---|---|---|
| **Post-dauer (PD)** | | |
| Dauer | + | - |
| Early PDL3 | ++ | - |
| Mid PDL3 | + | - |
| Late PDL3 | +++ | - |
| PDL3 molt | +++ | - |
| Early PDL4 | ++ | - |
| Mid PDL4 | ++ | - |
| Late PDL4 | ++ | - |
| Young PD adult | + | - |
| **Continuous** | | |
| Early L3 | ++ | ++ |
| Mid L3 | ++ | ++ |
| Late L3 | +++ | ++ |
| L3 molt | ++++ | +++ |
| Early L4 | ++++ | ++++ |
| Mid L4 | +++ | ++ |
| Late L4 | ++ | ++ |
| L4 molt | + | + |
| Young adult | + | + |

**Table S5. Oscillatory expression of a transcriptional reporter of *ztf-16.*** The *sIs12144[ztf-16p::gfp]* transcriptional reporter was observed during post-dauer and continuous development. Because expression was dynamic, larval stages were categorized as early, mid, or late, based on the extent of gonad and vulva development. At least 20 worms were observed at each stage.

| Relevant Genotype | % Adults Bursting |
|---|---|
| Wild-type | 0 |
| *let-7(n2853)* | 97 |
| *let-7(n2853) ztf-16(tm2127)* | 0 |
| *let-7(n2853) ztf-16(RNAi)* | 9 |

**Table S6. Reduced *ztf-16* activity suppresses the *let-7* bursting phenotype.** *let-7(n2853)* mutants burst at restrictive temperatures (24˚C), resulting in lethality. During continuous development, the loss or reduction of *ztf-16* through mutations or RNAi suppressed the bursting phenotype of *let-7(2853)* mutants. (n > 100 worms/strain).
